# Ecological and microbiological diversity of chigger mites, including vectors of scrub typhus, on small mammals across stratified habitats in Thailand

**DOI:** 10.1101/523845

**Authors:** Kittipong Chaisiri, A. Christina Gill, Alexandr A. Stekolnikov, Soawapak Hinjoy, John W. McGarry, Alistair C. Darby, Serge Morand, Benjamin L. Makepeace

## Abstract

Scrub typhus, caused by a bacterial pathogen (*Orientia* spp.), is a potentially life-threatening febrile illness widely distributed in the Asia-Pacific region and is emerging elsewhere. The infection is transmitted by the larval stage of trombiculid mites (“chiggers”) that often exhibit low host specificity. Here, we present an analysis of chigger ecology for 38 species sampled from 11 provinces of Thailand and microbiomes for eight widespread species. In total, >16 000 individual chiggers were collected from 1 574 small mammal specimens belonging to 18 species across four horizontally-stratified habitat types. Chigger species richness was positively associated with higher latitudes, dry seasonal conditions, and host maturity; but negatively associated with increased human land use. Human scrub typhus incidence was found to be positively correlated with chigger species richness. The bacterial microbiome of chiggers was highly diverse, with *Sphingobium, Mycobacterium, Neisseriaceae* and various *Bacillales* representing the most abundant taxa. Only *Leptotrombidium deliense* was found to be infected with *Orientia*. β-diversity, but not α-diversity, was significantly different between chigger species and geographic regions, although not between habitat types. This first field survey of the chigger microbiome provides a framework for future studies on interactions between pathogens and other symbionts in these understudied vectors.

## Introduction

The Trombiculoidea is a superfamily of mites (Acari: Acariformes) with a unique mode of parasitism among medically-relevant arthropod vectors. The larval stage, colloquially known as chiggers or berry bugs, is ectoparasitic on vertebrates (or occasionally invertebrates). In contrast, the deutonymph and adult stages have an edaphic lifestyle and are free-living predators of arthropods or their eggs [1]. Chiggers are the exclusive biological vectors of scrub typhus, a potentially life-threatening febrile illness of humans that historically has been associated only with the Asia-Pacific region [2]. However, recently endemic scrub typhus has been reported from the Middle East [3] and South America [4], and local transmission is suspected in sub-Saharan Africa [5]. The main aetiological agent of the disease, *Orientia tsutsugamushi* (Rickettsiales: *Rickettsiaceae*), is a vertically-transmitted chigger symbiont [6].

The epidemiology of scrub typhus remains poorly understood, largely because chiggers are minute (typically <250 μm in length) and very challenging to identify and utilise for molecular characterisation and screening [7]. In particular, interactions between climatic and physical geography, wild vertebrate hosts, and human disturbance of the environment with chigger species richness and abundance, and how these variables impact on scrub typhus incidence, are largely unexplored in most endemic regions. Moreover, our understanding of the bacterial associates of chiggers is mainly restricted to *O. tsutsugamushi* and a very small number of other potential human pathogens, such as *Bartonella* spp. [8] and *Rickettsia* spp [9]. As many cases of epidemiological-relevant interactions between human pathogens and the microbiome of arthropod vectors have been reported, our ignorance regarding the chigger microbiome is of potential concern for disease control. Indeed, this was highlighted recently by a 16S rRNA amplicon survey of a colony of the scrub typhus vector *Leptotrombidium imphalum*, which revealed a hitherto unrecognised association between a novel member of the *Amoebophilaceae* and *O. tsutsugamushi* in adult female mites [10]. The completion of the *Leptotrombidium deliense* genome project also uncovered an intimate relationship between chiggers and soil bacteria and fungi, as genes for secondary metabolism have been acquired by lateral transfer from these microorganisms [11].

Among scrub typhus-endemic countries, Thailand has some of the highest incidence rates. The Thai Bureau of Epidemiology reported an increase in annual minimum incidence from 6.0 per 100 000 persons in 2003 to 17.1 per 100 000 in 2013 [2]. The role of the vector in this increase is unknown, but the higher prevalence of *O. tsutsugamushi* in small mammal chigger hosts from forested regions relative to areas with greater human disturbance implicates land use as a key factor in disease risk [12]. Here, we present an analysis of chigger distributions on small mammals across 11 provinces of Thailand, their associations with habitat types stratified by human disturbance, and the microbiomes of nine widely-distributed chigger species. We show that chigger species richness is influenced by mammalian host status, climatic factors and land use; whereas chigger species and geographic region, but not habitat type, significantly impact on the β-diversity of chigger microbiomes.

## Materials & Methods

For a more detailed description of the Materials and Methods, see Supplemental Materials and Methods.

### Trapping of small mammals and chigger collections

This study utilised chigger material collected previously for a taxonomic study in Thailand [13]. In brief, small mammals were trapped across 13 localities between 2008 – 2015, once each in the dry season and wet season. Chiggers were removed from mammal cadavers and fixed in 70 - 95% ethanol. Mites collected from each animal were counted to estimate infestation intensity and chigger abundance, as defined by Rózsa *et al*. [14]. For identification and species richness estimation, 10-20% of chiggers from each host animal were selected using size and microscopic appearance as a guide to obtain a representative sub-sample.

### Ecological analysis

For ecological analysis, trapping sites were divided equally into four different types of habitats with respect to human land use (anthropization index), spanning low to high levels of disturbance [15–17]. Calculation of infestation intensity statistics of conspecific chigger species were performed using the “BiodiversityR” package. In addition, 12 chigger species that infested ≥10 individual hosts were included in an analysis of association with habitat type using the “FactoMineR” package in R.

### Network analyses of host-chigger interactions

To study the community ecology of host-chigger interactions, bipartite network analyses of host-ectoparasite interactions were conducted at both community (pooled host species or pooled locations) and individual levels using “vegan” [18] and “bipartite” packages [19] implemented in R freeware. Bipartite networks were transformed to unipartite networks using the “tnet” package [20]. Unipartite network plots illustrate relative interaction patterns within a host community with respect to the co-occurrence of chigger species.

### Multiple regression models of independent variables explaining chigger species richness

Generalized linear models were constructed in order to identify potential effects of host attributes (species, sex, maturity and body mass) and ecological factors (habitat, site and season) on chigger species richness at the individual host level. Poisson regression models were created for chigger species richness count data using the “lme4” package [21] in R freeware. Selection of models was based on Akaike’s Information Criterion adjusted for small sample size (AICc) using the “gmulti” package [22] in R freeware. Data for scrub typhus human case numbers from the 13 studied sites were obtained from the Bureau of Epidemiology, Ministry of Public Health, Thailand (unpublished data).

### DNA extraction

As clearing in Berlese’s fluid destroys DNA, chiggers destined for DNA extraction were identified using autofluorescence microscopy as previously described [7]. Genomic DNA was purified using the DNeasy Blood & Tissue Kit (Qiagen, Hilden, Germany).

### Library preparation and next generation sequencing of 16S rRNA amplicons

To determine the bacterial microbiome of chiggers, a dual-index nested PCR protocol for MiSeq (Illumina, San Diego, CA, USA) sequencing was applied [23–25] targeting the v4 region of the 16S rRNA gene. The second round indexing PCR was performed using the Nextera XT DNA protocol (Illumina). Each MiSeq run included three types of negative control to identify potential background contamination from sample manipulation equipment, DNA extraction kits and PCR reagents used in library preparation. Samples were submitted for sequencing with 300 bp paired-end chemistry on the Illumina MiSeq platform at the Centre for Genomic Research (University of Liverpool).

### Microbiome profiling

Analyses of 16S rRNA microbiome profile were performed using the Quantitative Insights into Microbial Ecology (QIIME) software package, version 1.8.0 [26]. Operational taxonomic units (OTUs) were created using an open-reference approach using the USEARCH61 method [27] whereby reads are binned at 97% similarity [27] against the Greengene database v. 13_8 [28] followed by *de novo* OTU picking. Bacterial taxonomic assignment was performed with UCLUST. Chimeric sequences were removed using “ChimeraSlayer” [29].

### Comparative analyses of the chigger microbiome

Read counts were normalized to relative abundance for graphing or rarefied to 10 000 reads for diversity calculations. Bacterial communities were categorised according to sample type (individuals and pools), selected chigger species and study sites (mixed species), as well as soil samples from Thailand and Lao PDR. For details of α- and β-diversity analyses, and principal coordinates analysis (PCoA), see Supplemental Materials and Methods.

### Geobacillus *qPCR and Sanger sequencing*

A pair of PCR primers was designed to amplify a 16S rRNA gene portion for the genus *Geobacillus* and related Firmicutes. Individual, 25-pooled and 50-pooled chiggers, as well as water samples from the laboratory water bath (Grant Sub; Grant Instruments, Cambridge, UK) and Qiagen microbial DNA-free water (negative control), were used in the qPCR assay. DNA from chiggers and 10 μl of water bath samples were extracted using the DNeasy Blood & Tissue Kit (Qiagen).

Bacterial taxonomy was assigned using RDP Classifier Version 2.10 [30] available at https://rdp.cme.msu.edu, using a >80% confidence threshold [31]. The DNA sequences were aligned using ClustalW and phylogenetic tree construction was performed with the maximum likelihood method using Mega software version 6.06 [32].

### Determination of GC content in 16S rRNA sequences

We evaluated whether the influence of GC content differentially affected data obtained from individual and pooled chiggers (low and high DNA concentration templates, respectively). Representative sequences of the dominant bacterial OTUs from individual and pooled chiggers were assessed for GC content using “Oligo Calc”, an oligonucleotide properties calculator available at http://biotools.nubic.northwestern.edu/OligoCalc.html [33] and their mean GC content was compared by two-sample *t*-test.

## Results

### Chigger ecology and host associations

A total of 16 761 chiggers were obtained from 1 574 small mammals belonging to 18 species. The overall infestation rate was 23.8%, with Bo Kleu district (Nan province) displaying the highest rate recorded for a single site (95%) (Table S1). The highest mean chigger intensity (113.3) was observed in *Berylmys bowersi* (Bower’s white-toothed rat) (Table S2). A subsample of 2 519 chiggers (approximately 15% of the total) were identified to the species level, revealing that *Rattus tanezumi* (Asian house rat) and *Bandicota indica* (greater bandicoot rat) exhibited the greatest chigger species richness (21 species each). Approximately half of the infested hosts (50.7%) harboured a single chigger species, 33.3% harboured two, and the remainder harboured ≥3 species. *Ascoshoengastia indica* was most prevalent (7.31%; the only species recorded from every geographic region), followed by *L. deliense* (5.22%) and *Walchia micropelta* (5.16%) (Table S3).

A species accumulation curve plot demonstrated that the sample size of small mammals was sufficient to describe the chigger species diversity accurately, since a plateau was reached at around 1,000 hosts (Fig. S1). Chigger species richness of sampling locations increased at higher latitudes (Spearman’s rank correlation = 60.81, p = 0.0023; Fig. S2) and varied significantly among the four habitat types (in descending order) of forest, dry land, rain-fed land, and human settlement at both an individual host level (Kruskal-Wallis statistic = 91.29, df = 3, p < 0.0001; Fig. 1b) and for the whole population (Fig. 1a). Moreover, while there were no seasonal differences in chigger species richness or abundance at the individual host level, chigger species richness was considerably higher in the dry season than in the wet season at the whole country level (Fig. 1c). Ecological specialisation of some of the most widespread chigger species (*A. indica, W. micropelta* and *Walchia pingue*) between habitat types was weak (Fig. 2). However, *L. deliense* showed a preference for areas in forest or dry land; whereas other species with more restricted distributions displayed predilections for human settlements (*Helenicula kohlsi*), rain-fed lowland (e.g., *Walchia minuscuta*) or dry landscapes (*Helenicula pilosa*) (Fig. 2).

**Figure 1:**
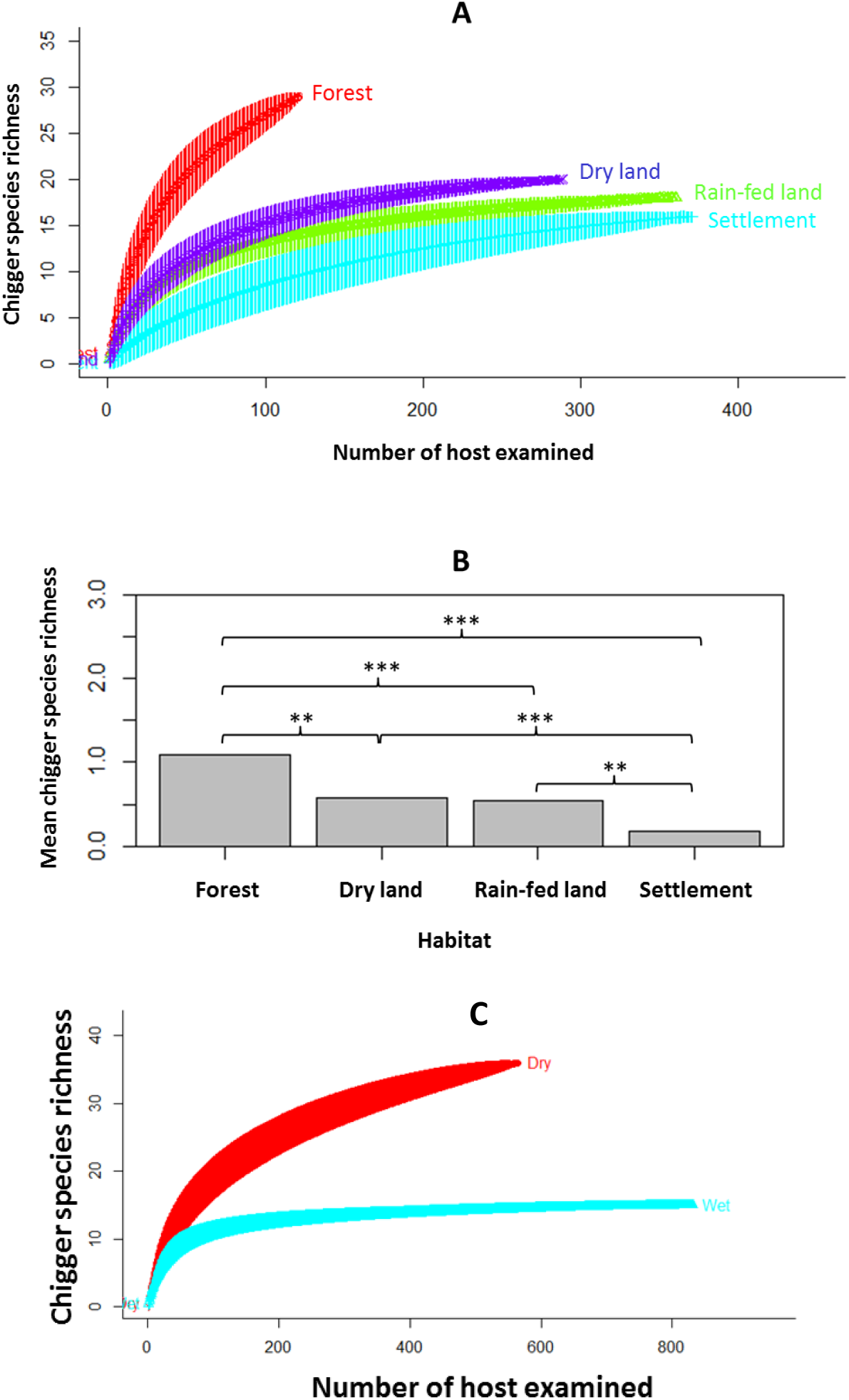
Effect of habitat and season on chigger species richness. (A) Chigger species accumulation curves among different habitats at the host population level. (B) Mean chigger species richness per host individual by habitat type. (C) Chigger species accumulation curves between the dry (red) and wet (blue) season.

**Figure 2.**
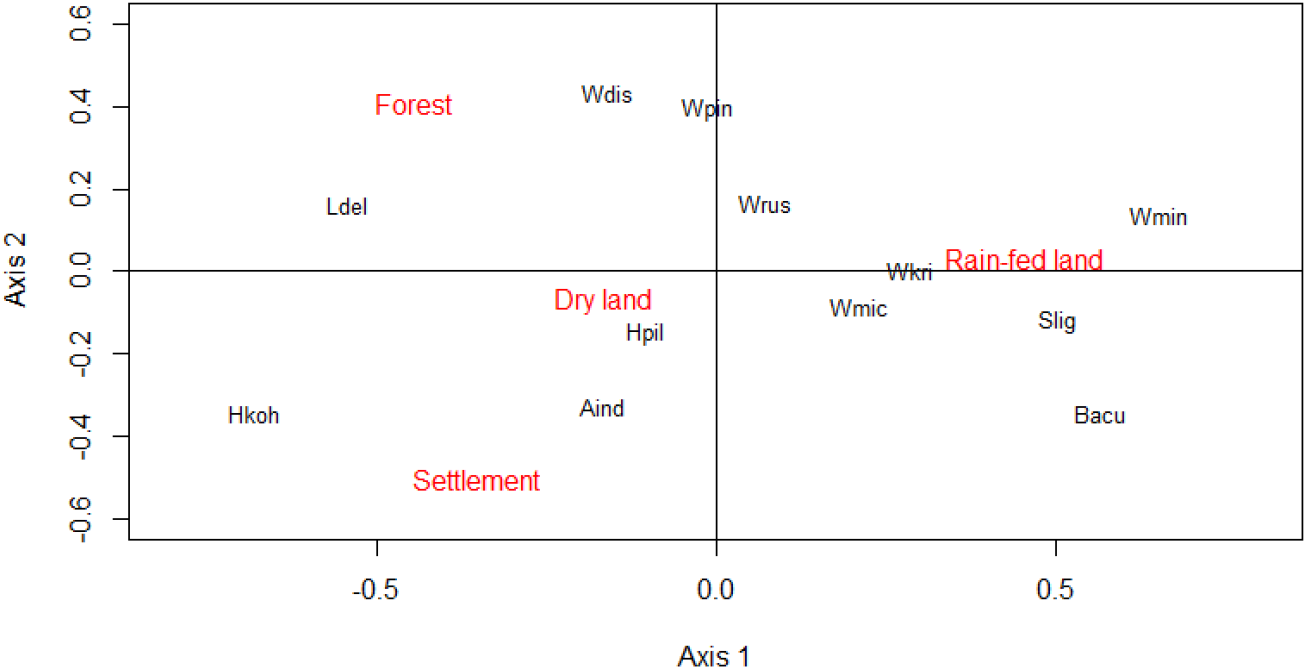
Correspondence analysis showing the association between the 12 dominant chigger species (Aind, *Ascoschoengastia indica*; Bacu, *Blankaartia acuscutellaris*; Hkoh, *Helenicula kohlsi*; Hpil, *Helenicula pilosa*; Ldel, *Leptotrombidium deliense*; Slig, *Schoengastiella ligula*; Wdis, *Walchia dismina*; Wkri, *Walchia kritochaeta*; Wmic, *Walchia micropelta*; Wmin, *Walchia minuscuta*; Wpin, *Walchia pingue*; Wrus, *Walchia rustica*) within the four categorized habitats. The first and second dimensions explain 87% of the total variance (axis 1, 59.82%; axis 2, 27.38%).

Bipartite network analysis showed highly complex interactions between chigger and host species (Fig. 3a). The largest chigger species assemblages at the whole host population level were found on two rodent species associated with human settlements, *B. indica* and *R. tanezumi*. Interestingly, the only non-rodent hosts sampled in this study, *Hylomys suillus* (Erinaceomorpha: Erinaceidae) and *Tupaia glis* (Scandentia: Tupaiidae), were parasitized by several chigger species never found on rodents (Fig. 3a). Overall however, more than half of chigger species were found on more than one host species, and the species-specificity for those found on >10 individual animals was only 0.171 – 0.542. A unipartite network analysis supported the bipartite analysis, assigning *B. indica* and *R. tanezumi* with the highest Eigenvector centrality scores among all of the hosts (Fig. 3b).

**Figure 3.**
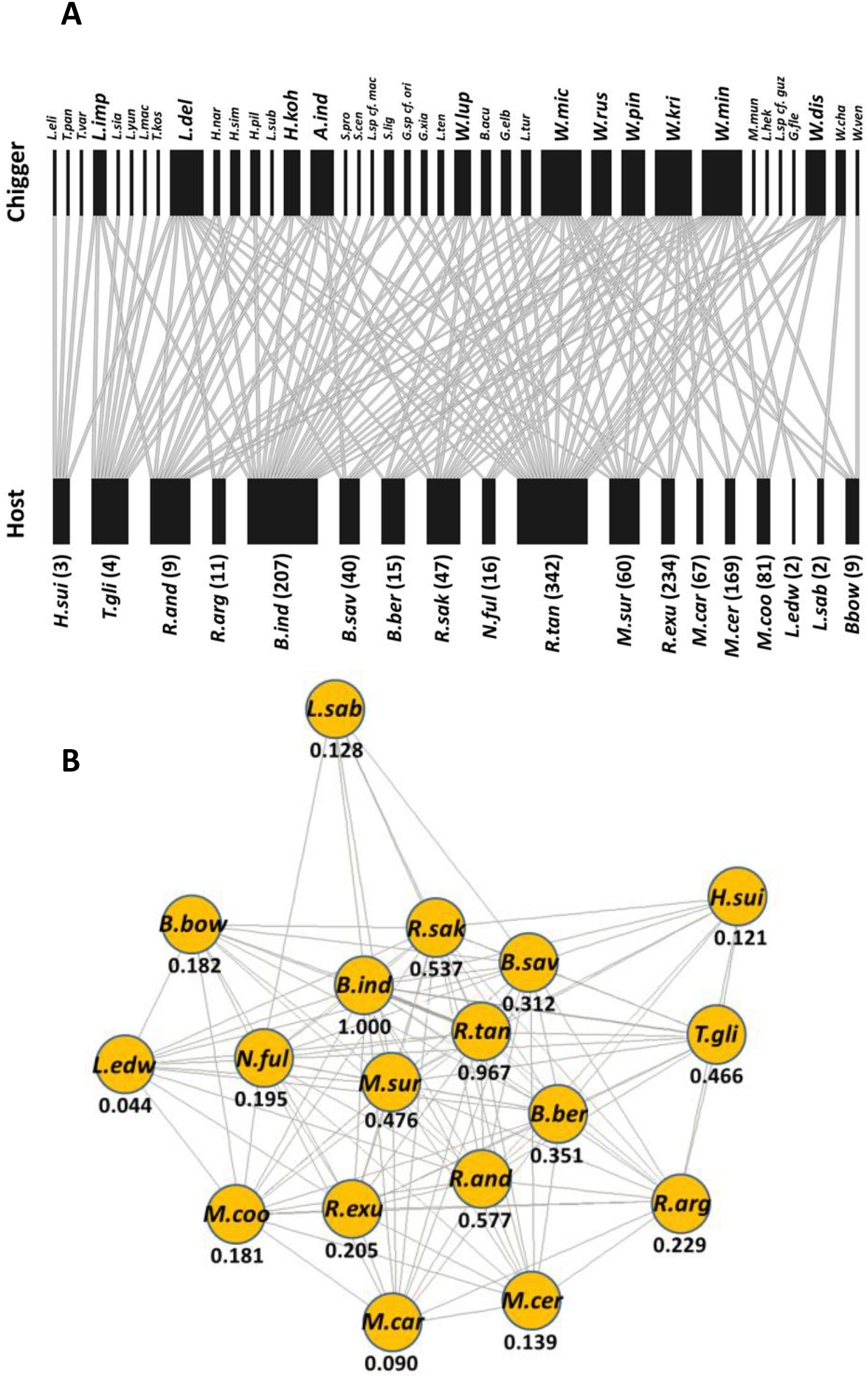
Host-chigger associations in Thailand. (A) Bipartite graph based on presence-absence data. The number of individual hosts examined is shown in parentheses. Chigger species with broad host ranges are displayed in bold. (B) Unipartite network and Eigenvector centrality scores illustrating the pattern of chigger sharing among 18 small mammal hosts.

### Chigger-host properties and scrub typhus incidence

For each of the 13 geographic sites, bipartite network properties of host-chigger interactions were calculated at the individual host level, including the nestedness metric based on overlap and decreasing fill (NODF), network connectance, links per species, and network modularity. The highest NODF and connectance were found in the Nakhonsawan network, where chigger species richness was only four species; while the Chiangrai network exhibited an elevated chigger species richness (12 species), but with the lowest NODF and connectance (Table S4). In contrast, Chiangrai displayed the highest modularity within the network, whereas the least network modularity was found in Prachuab Kirikhan (Table S4).

We tested the effect of various independent variables on individual chigger species richness using GLMs with model selection by Akaike’s Information Criterion. Host species, host maturity, site and habitat (but not host sex) were significant variables in the best 10 models (Table S5; Fig. S3a). Animals captured in forest demonstrated significantly higher chigger species richness than hosts from human settlements (estimate = −1.074, p <0.0001; Table S6), and species richness was greater on mature hosts than on juveniles (estimate = −0.283, p = 0.004; Table S6).

We then applied the same modelling approach but included human scrub typhus cases at district level with environmental variables (elevation, annual mean temperature and latitude; Table S7), chigger species richness, and network properties (Fig. S3b). Network connectance and chigger species richness strongly influenced local scrub typhus case numbers, as the two variables appeared in the top 10 selected models (Table S8). Finally, we performed a univariate analysis, which also showed that scrub typhus case number was positively correlated with chigger species richness (Spearman rank correlation = 45.71, p = 0.0006; Fig. 4a) and negatively correlated with host-chigger network connectance (Spearman’s rank correlation = 485.45, p = 0.011; Fig. 4b). Importantly, there was no significant relationship between overall chigger abundance and scrub typhus incidence (R^2^ = 0.105, *P* = 0.732; data not shown).

**Figure 4.**
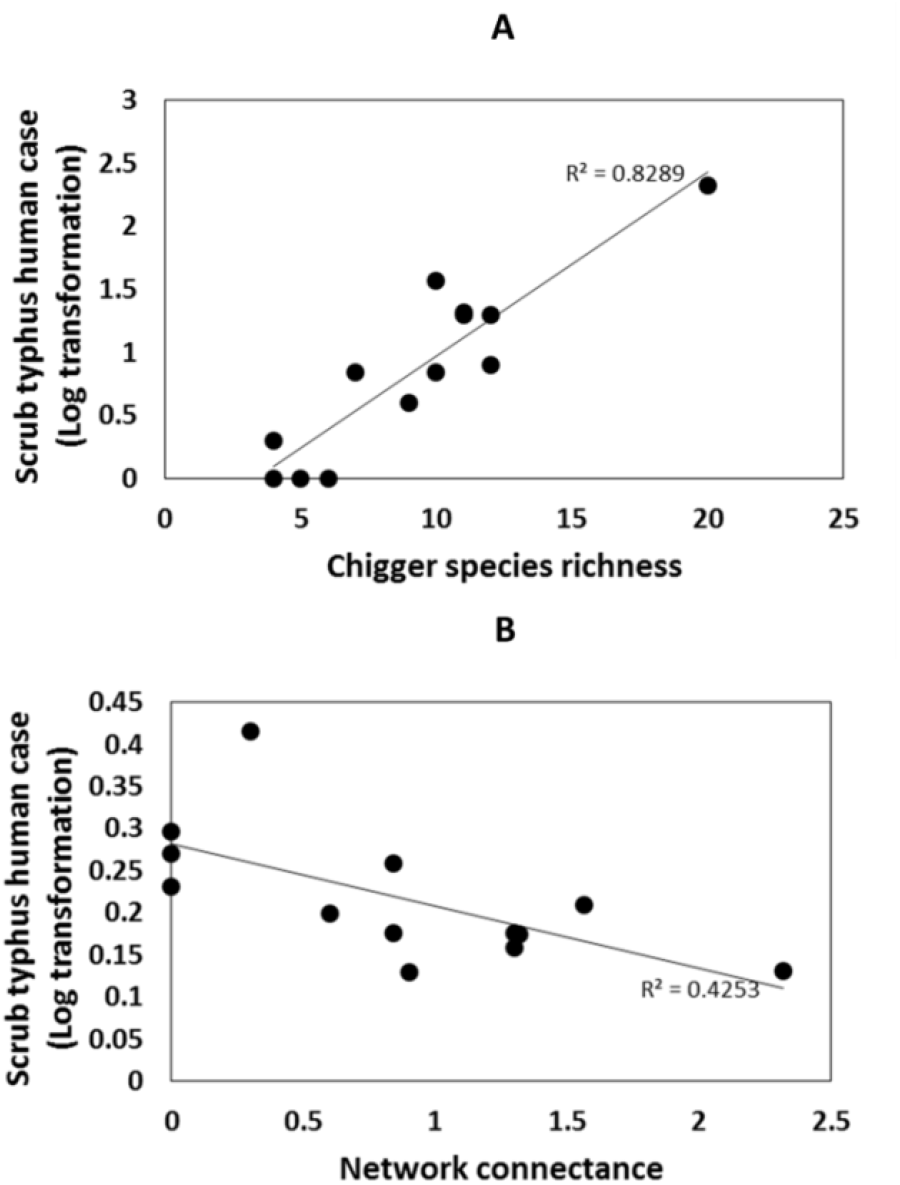
Correlation plots show the relationship between chigger ecology [(A) chigger species richness; (B) host-chigger network connectance] and scrub typhus incidence in humans. Incidence data are displayed as the log_10_ transformation of the number of cases per year.

### Microbiome of individual and pooled chigger specimens

The total number of 16S rRNA reads from the complete set of 366 samples (264 individual chiggers, 69 pooled chiggers, 18 soil samples and 15 background controls) after quality filtering, de-multiplexing and error correction was 51 896 654 (mean reads per sample = 137 657; SD = 69 522). After paired read alignment and size selection at 270 - 300 bp, the read number was 49 635 427 (mean reads per sample = 131 659; SD = 69 922), a sequence retention of 94%. The analysis of individual chigger specimens comprised nine widespread species: *A. indica, L. deliense, W. micropelta, W. minuscuta, Walchia kritochaeta, H. pilosa, H. kohlsi, Blankaartia acuscutellaris*, and *Schoengastiella ligula*. However, after removing samples with high similarity to negative controls (see Supplemental Materials and Methods), more than half (58.7%) were excluded from downstream analyses, including all of those for *W. minuscuta*.

The microbiomes of individual chiggers were dominated by several different *Geobacillus* OTUs (Fig. 5). However, *Sphingobium* (α-Proteobacteria) was also abundant, as were Comamonadaceae (especially in *Walchia* spp.) and *Brevibacillus* (particularly in *B. acuscutellaris* and *L. deliense*). Importantly, we only detected *O. tsutsugamushi* in *L. deliense* (3/39 individual specimens that passed QC), with a maximum OTU proportion of 19.58% (Fig. 5; Table 1). Other bacteria with pathogenic potential in humans were found across several chigger species, including *Mycobacterium* (11.93% of specimens), *Staphylococcus* (8.25%) and *Haemophilus parainfluenzae* (7.34%) (Table 1). However, most arthropod symbionts known to be of importance in other mite species or in insects (*Cardinium, Pseudonocardia* and *Rickettsiella*) were rare (<2% prevalence), while *Wolbachia* remained undetected at the individual level (Table 1).

**Figure 5.**
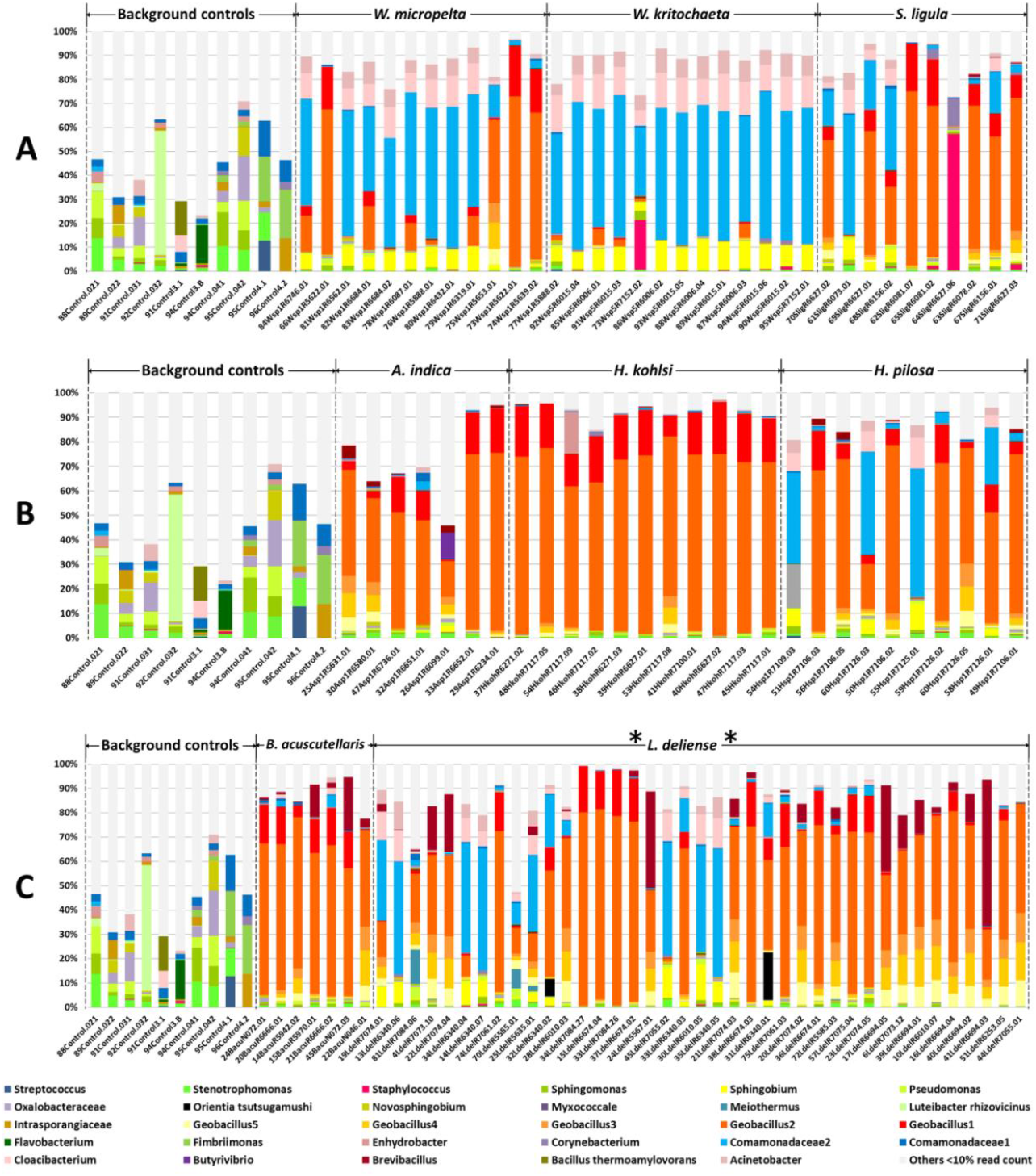
Relative abundance of bacterial OTUs in background controls and individual chiggers. (A) subfamily Gahrliepiinae and subfamily Trombiculinae. (B) Tribe Schoengastiini. (C) Tribe Trombiculini. The data are filtered; OTUs that represented <10% in a sample were combined in “others” (light grey) to aid visualisation. Source data are included in Table S11.

**Table 1.**
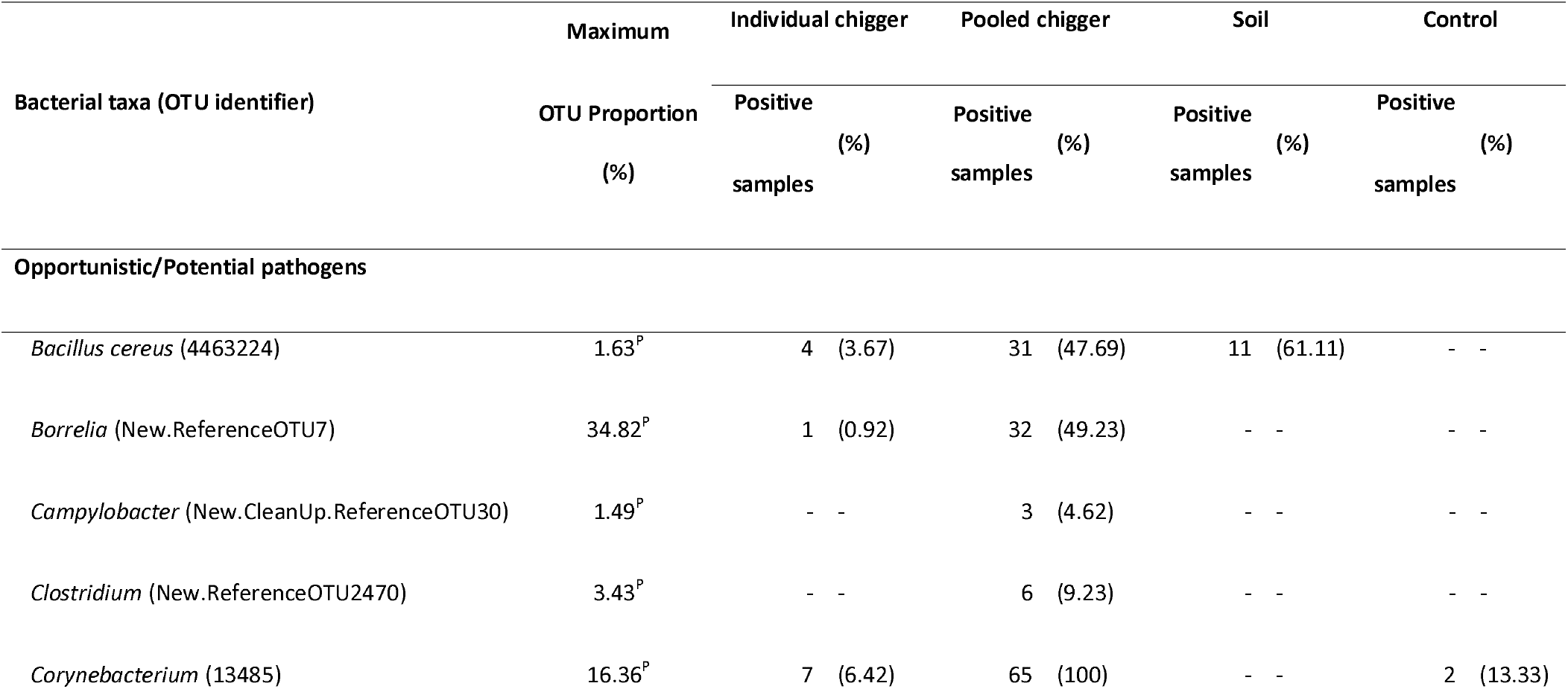
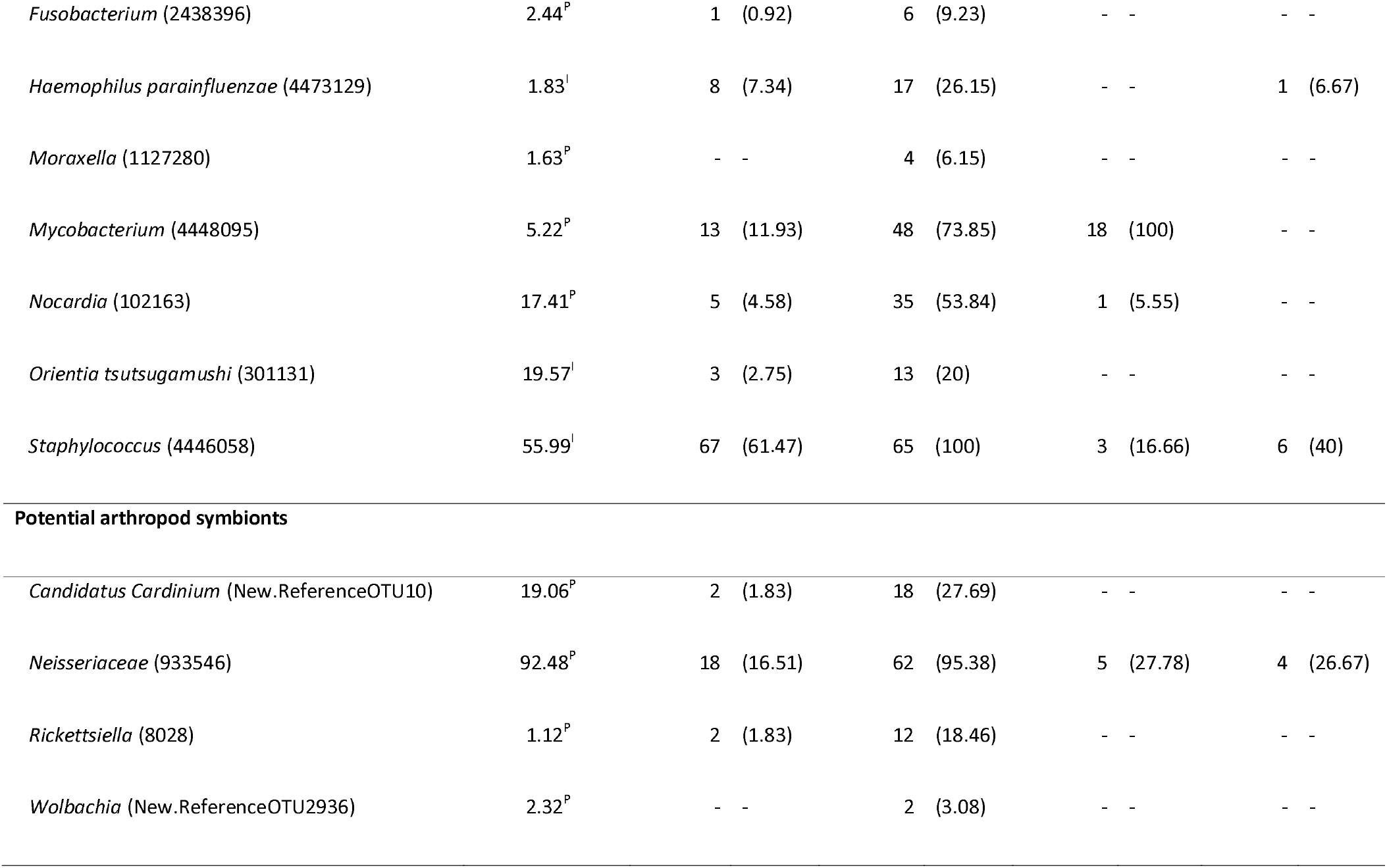
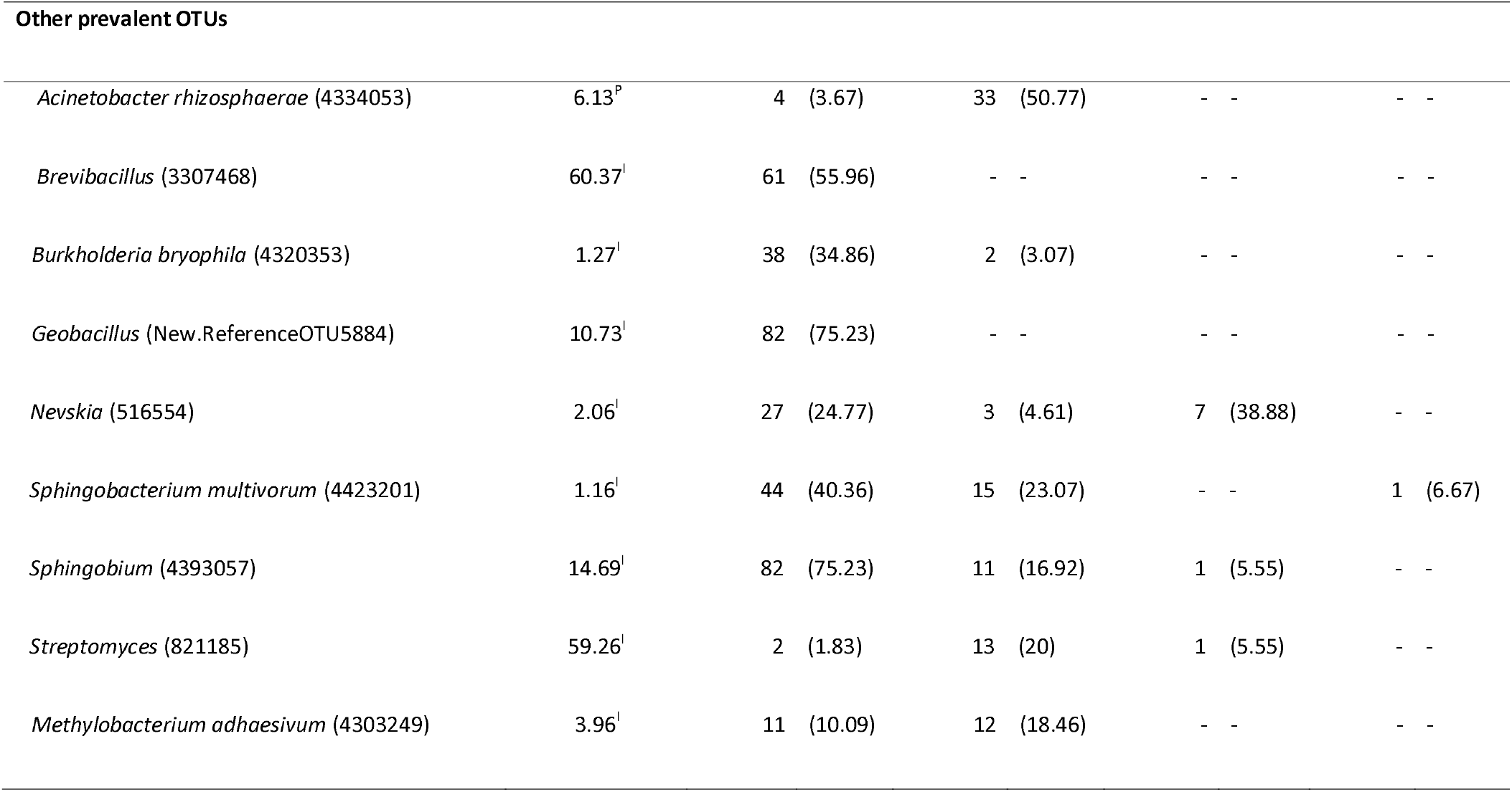
Selected bacterial taxa of public health importance, potential symbionts, and other prevalent OTUs detected in individual and pooled chiggers in comparison to soils and background controls. Only the OTUs with ≥5 reads were included. Superscript “I” or “P” indicate maximum OTU proportion values from individual or pooled chiggers, respectively.

To mitigate the problem of low biomass when amplifying 16S rRNA fragments from individual chiggers, we also sequenced several pools of 50 specimens each for *A. indica, L. deliense, W. micropelta, W. minuscuta*, and *Blankaartia acuscutellaris* (Fig. 6a); as well as three mixed-species pools of 50 specimens each for all 11 Thai provinces (Fig. 6b). This strategy was successful, as fewer samples (7.2%) were removed due to high similarity with negative controls when compared with individual samples. Surprisingly, two OTUs (*Geobacillus* and *Brevibacillus*) that were highly prevalent and relatively abundant in the individual-level data were not present at a read count ≥5 in any of the pooled data (Table 1). For some of the potential pathogens, individual and pooled data showed good concordance (*Staphylococcus* and *Mycobacterium* detected in 95.38% and 73.85% of pools, respectively), whereas others that were rarely detected in individuals were robustly confirmed by the pooling strategy (*Borrelia* in 49.23% and *Corynebacterium* in 78.46% of pools, respectively) (Table 1). Pooling also provided additional evidence that three classical arthropod symbionts (*Cardinium, Pseudonocardia* and *Rickettsiella*) were present in chiggers (~20 – 45% of pools), while a fourth (*Wolbachia*) was present only in two (3.08%) pools (Table 1). Intriguingly, one *Neisseriaceae* OTU (933546) was detected in 95.38% of pooled samples and was particularly dominant in *L. deliense*, reaching a maximum OTU proportion of 92.48% (Table 1). In accordance with the individual chigger data, the 13 (20%) pooled samples positive for *O. tsutsugamushi* (Table 1) all contained *L. deliense*.

**Figure 6.**
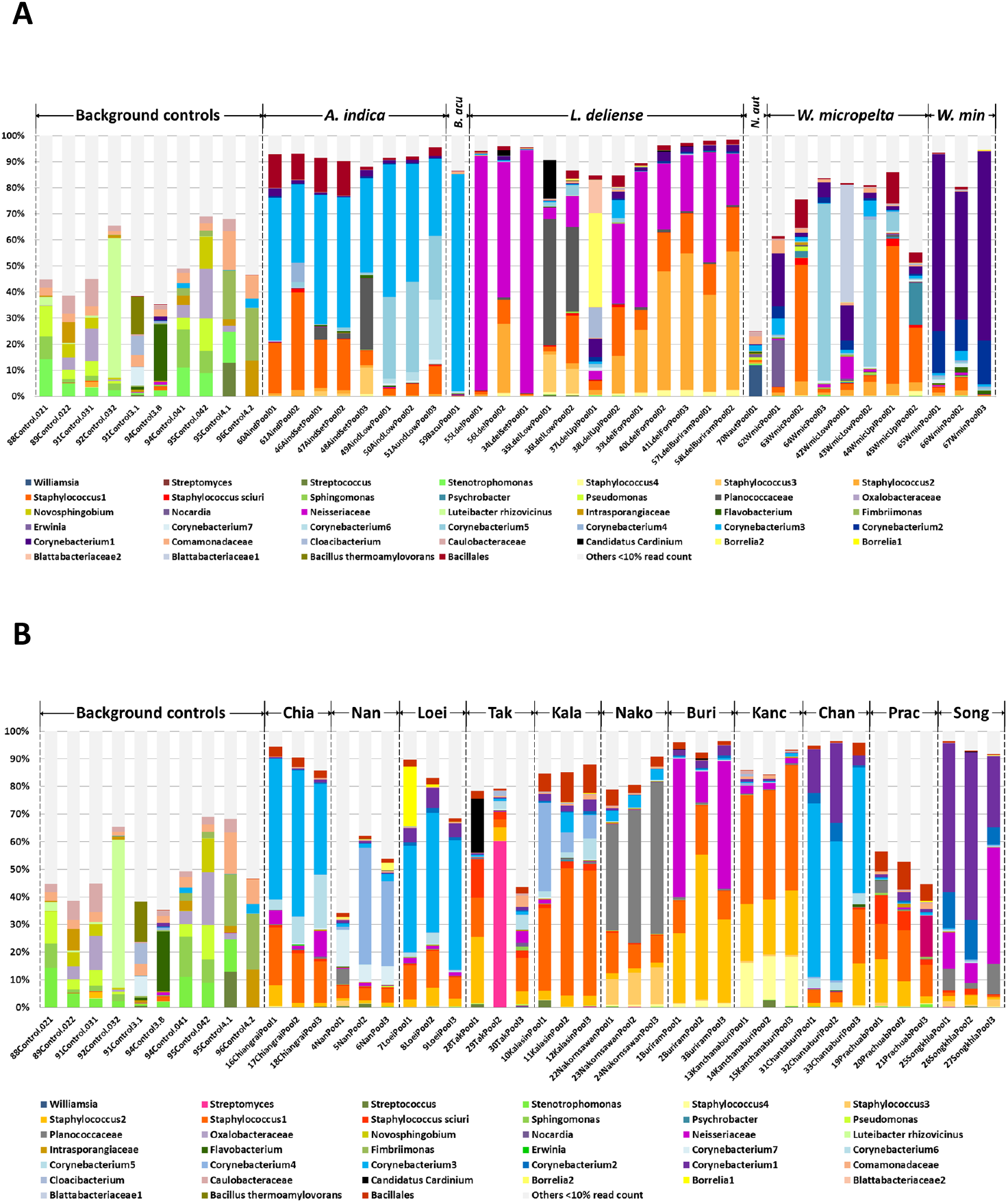
Relative abundance of bacterial OTUs in background controls and pooled samples. (A) Pools by chigger species (50 individuals per sample). (B) Mixed chigger species (50 individuals per sample) pooled by province. The data are filtered; OTUs that represented <10% in a sample were combined in “others” (light grey) to aid visualisation. Source data are included in Table S11

To investigate whether the presence of *Geobacillus* might have resulted from contamination of samples with spores in the laboratory or with bacterial DNA in the extraction kits, we first examined OTUs sequenced from negative controls, then measured levels of Firmicutes 16S rDNA by qPCR in chiggers compared with samples from the laboratory water bath. The dominant *Geobacillus* OTU observed in individual chiggers was absent in background controls (Table 1; Table S9). Despite a high Firmicutes signal in the water bath (Fig. S4), Sanger sequencing revealed that this was derived from *Paenibacillus* spp. and related *Bacillales*, whereas *Geobacillus* spp. were only observed in individual chigger samples (Fig. S5). Finally, we calculated percent GC content for the 15 most abundant OTUs in individual specimens and the 26 most abundant OTUs in pooled samples. This showed that the GC content of the OTUs from individual specimens was significantly higher than for the pooled material (*P* = 0.0094; Fig. S6).

### Factors affecting the microbial profile of chiggers

The α-diversity of bacterial OTUs determined by a richness estimator (Chao1) and the whole-tree phylogenetic diversity index (PD_whole_tree) revealed significant differences between the sample types, with pooled chigger and soil samples exhibiting higher diversity than individual chigger specimens (Kruskal-Wallis test with post-hoc Bonferroni correction, *P* <0.001) (Table S10). The latter were not significantly more diverse than control samples. Analysis of β-diversity showed that the sample types were generally well separated from one another (ANOSIM: *R* = 0.7997, *P* = 0.001), although some background controls were nested on the periphery of the individual chigger samples (Fig. S7). Bacterial communities were significantly clustered with respect to chigger species and geographic location (study sites) in both individual and pooled chiggers (*P* <0.001), whereas habitat (human disturbance transect) failed to show a significant effect (Fig. 7). The impact of chigger species and geographic location on β-diversity displayed similar correlation coefficients and network topology (Fig. 7).

**Figure 7.**
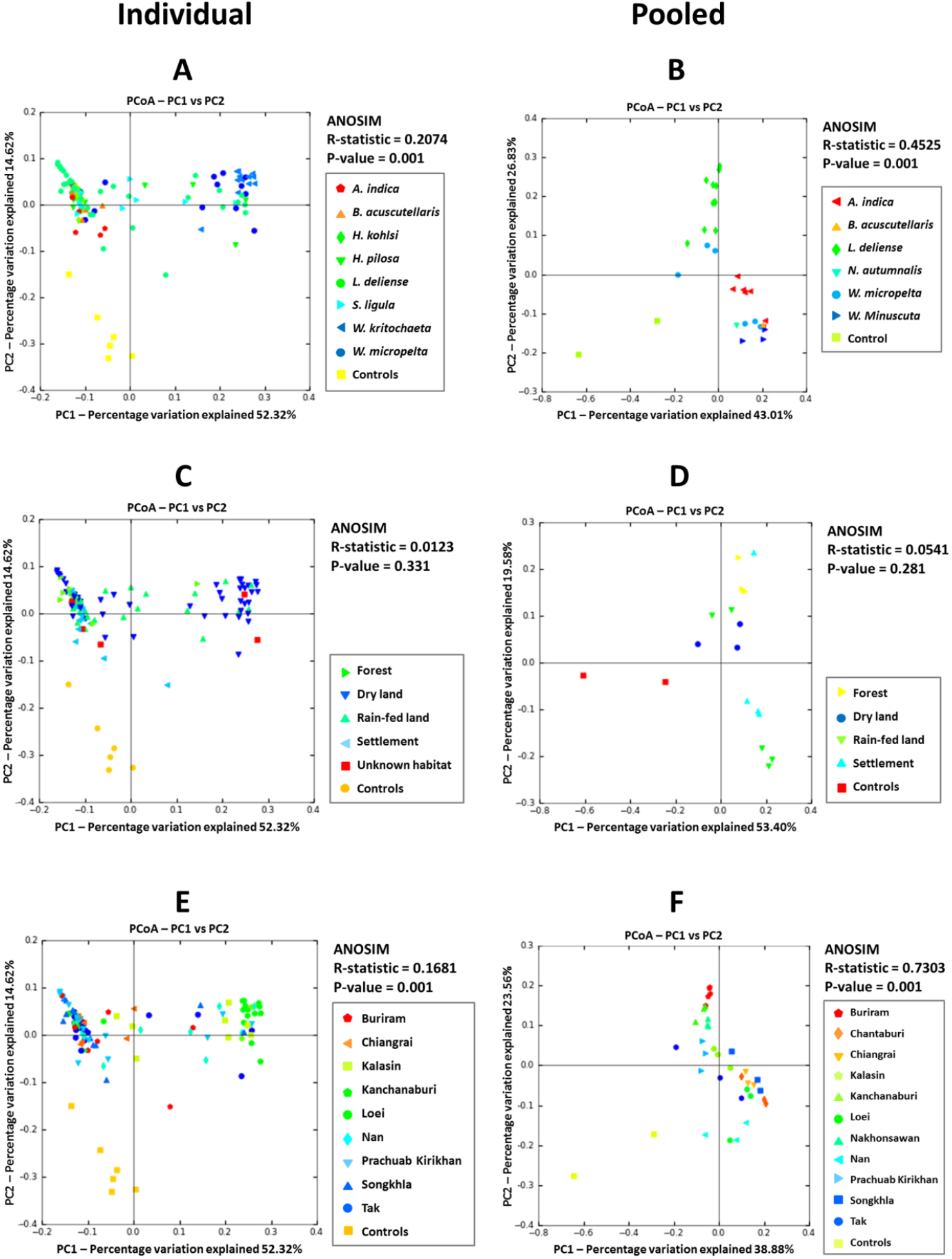
Principal coordinates analysis plots created using the unweighted UniFrac metric showing bacterial community clustering of individual (left panels) and pooled chiggers (right panels) among different sample categories. (A, B) chigger species; (C, D) habitat; and (E, F) study site. Control data are shown for reference only and were not included in the ANOSIM.

## Discussion

To the best of our knowledge, habitat type, chigger diversity and human scrub typhus incidence have never been analysed together on a countrywide scale before. The current study sampled more than one-third of known chigger species in Thailand [13] and found three significant environmental associations with species richness: a positive correlation with latitude, a negative correlation with increased human land use, and an elevated level of chigger diversity in the dry season (in addition to significant effects of host species and maturity). Latitudinal gradients are associated with bioclimatic factors such as mean temperature, humidity and rainfall, and the species richness of animals and plants tends to increase at lower latitudes approaching the equator [34–36]. Thus, parasite diversity might also be expected to be higher at lower latitudes, and there is some evidence of this for microbial pathogens [37–39]. However, the opposite trend observed here is supported by previous studies on fleas [40]. Moreover, in Yunnan province in China, chigger diversity on small mammals was even higher than in our study and increased according to latitude up to a zenith at 25 - 26°N before decreasing further north [41], suggesting the presence of an optimum zone (our study covered 7 – 19°N). One hypothesis to explain this phenomenon is that the geographic range of individual hosts tends to be broader at higher latitudes, perhaps facilitating the accumulation of a greater diversity of ectoparasites [36].

Analysis of chigger distributions in Yunnan also concur with our findings on the impact of human disturbance of the natural environment, with a significantly greater host and chigger species richness being observed in a mountainous uncultivated landscape habitat compared with a cultivated flatland landscape [42]. We increased the resolution of the human land use analysis in our study by trapping hosts across a transect of four rather than two habitat categories, which revealed a stepwise reduction in chigger species richness as human disturbance increased, reflecting the universal process of loss of animal and plant diversity through urbanisation. Seasonality was also an apparent determinant of chigger diversity in our study, with a striking increase in species richness during the dry season compared with the wet season. However, caution is required in interpreting this finding, as our field studies were not designed to standardise sampling across the two seasons. Nevertheless, it is plausible that breeding frequency is reduced in the wet season, and/or that many chiggers emerging from underground during monsoon periods are washed into water bodies before they can attach to a host. Chigger species also have different seasonal preferences. For instance, in subtropical regions, most scrub typhus cases occur in the autumn when populations of *Leptotrombidium pallidum* and *Leptotrombidium scutellare* dramatically increase (as seen in South Korea [43, 44]), or the main vector may change between summer and winter (as seen with *L. deliense* and *L. scutellare* in Taiwan, respectively [45]).

Here, the species richness of chiggers was identified as a positive correlate of scrub typhus incidence for the first time. Since decreased human land use is associated with both increased chigger species richness (this study) and higher prevalence of *O. tsutsugamushi* infection in small mammals [12], increased biodiversity may be a risk factor for human scrub typhus. This contradicts a meta-analysis of chigger-host relationships in Yunnan, where a lower host and chigger diversity in cultivated flatland was associated with a greater abundance of chiggers, especially species that are known or potential vectors of scrub typhus [42]. However, as neither human scrub typhus incidence nor *O. tsutsugamushi* prevalence in small mammals was incorporated into the Yunnan study, and an efficient scrub typhus vector (*L. scutellare*) was abundant in mountainous uncultivated land (the higher biodiversity site), the impact of land use on infection risk remains an open question in that region. In Taiwan, spatial modelling of land use data revealed significant positive correlations between crop-vegetation mosaics and forest, as well as elevation, with scrub typhus incidence [46]. In contrast with our study, follow-up investigations in Taiwan found that chigger prevalence and abundance on small mammals was positively associated with both human scrub typhus incidence and host *O. tsutsugamushi* seropositivity [45]. Chigger species richness and host-chigger networks were not explicitly incorporated in this Taiwanese study, but both the diversity of chiggers (12 species) and their hosts (8 species) were markedly lower than we observed in Thailand.

At first glance, the association between chigger species richness and scrub typhus incidence in Thailand seems paradoxical as we only detected *O. tsutsugamushi* in a single species (*L. deliense*). Yet it is important to emphasise that other potential vectors of scrub typhus (*e.g., L. imphalum*, the principal vector in northern Thailand [47]) were collected but not subjected to 16S rRNA sequencing. Moreover, more than 20 species of *Leptotrombidium* have been reported from Thailand, many only from the northern provinces where scrub typhus incidence is highest [13]. Although the majority of these species are not known to be scrub typhus vectors, recent data on vector competence are lacking, and it is plausible that transmission of *O. tsutsugamushi* by two or more vectors in the same region may contribute to diversification of the pathogen and an increase in human cases, as has been recently hypothesised for Taiwan [45, 48]. We also observed that human scrub typhus incidence was negatively associated with host-parasite network connectance, suggesting that increasing complexity of chigger-host interactions might reduce human exposure by zooprophylaxis, or lead to a greater likelihood that non-vector species dominate networks.

With the exception of a laboratory colony of *L. imphalum* [10], the composition of the chigger microbiome was largely unknown prior to our study. Our data reveal complex microbiomes that (in contrast with those of many other arthropods such as certain vectors [49] or sap-feeding insects [50]) are not dominated by a very small number of specialised primary and secondary symbionts. Since questing chiggers emerge from underground and are associated with their host for only few days before moulting into free-living nymphs, we hypothesised that they may not require symbionts for dietary supplementation, and may passively accumulate soil microbes instead. Indeed, it is now known that the *L. deliense* genome contains terpene synthase genes that appear to have been acquired by ancient lateral gene transfer from Actinobacteria and other environmental phyla [11]. However, while bacterial sequences of putative soil origin were prevalent in chiggers (*e.g., Bacillus cereus* and *Mycobacterium* spp.), on the basis of the limited number of soil samples we analysed here, the chigger “microbiome” is not simply a result of soil particles adhering to the mite surface. The clear impact of chigger species and geographical location, but not human land disturbance, on the microbial sequence profiles lends further support to the concept of an integral microbiome in chiggers that may be modulated by habitat on large (several hundred km) but not small (a few km) scales. This may be because their mobile hosts can travel between the human disturbance zones we defined within sampling sites [16, 51]. Evidence that the host also contributes to the chigger microbiome was revealed by the presence of typical mammalian-associated flora such as *Staphylococcus* spp. and *Haemophilus* spp.

The prevalence of the classical intracellular arthropod symbionts *Cardinium, Rickettsiella* and *Wolbachia* was quite low in individual specimens, despite their importance in other mite taxa [52]. While *Orientia* in some *Leptotrombidium* spp. could conceivably displace these symbionts due to competition for intracellular niches, it is rare or absent in most other chigger genera [53]. Unfortunately, the sample size of *Orientia*-infected chiggers was too small in this study to investigate the impact of the pathogen on microbiome composition. In contrast to a recent analysis of the microbiome of colonised *L. imphalum*, we found no evidence for an abundant *Amoebophilaceae* OTU; although this is not surprising, as it was found to be uncommon in all life stages except *Orientia*-infected adult females [10], and neither this species nor life stage was included in our microbiome analysis. Future studies should consider the role of *Neisseriaceae* OTU 933546 in chigger biology and potential interactions with vectored pathogens. Notably, this family in the β- proteobacteria exhibited a moderate prevalence in *L. deliense* individuals and contains gut symbionts of bees (*Snodgrassella alvi* [54]) and termites (*Stenoxybacter acetivorans* [55]). This suggests a facultative relationship in *L. deliense*, especially as OTU 933546 was also found in almost 30% of soil samples.

The high prevalence of sequences from *Geobacillus* spp. was surprising, since this is a thermophilic, spore-forming genus with an optimal growth range of 45 - 70°C. *Geobacillus* spp. thrive in hot composts, subterranean oilfields and hydrothermal vents, but due to their exceedingly robust spores that can be transported worldwide in atmospheric currents, isolates have been obtained across a vast range of temperate or cold terrestrial and marine sediments [56]. We did not surface-sterilise the chiggers as there is no evidence this procedure significantly affects microbiome data obtained from arthropods [57], and the risk of degradation of internal DNA when dealing with soft, minute species that might exhibit small breaches in their exoskeleton was acute. In any case, the dominant *Geobacillus* OTU detected in individual chiggers was absent from the soil we analysed. Despite the potential ubiquity of *Geobacillus* spp. spores in the environment, it is intriguing that this genus is not observed more frequently in arthropod microbiomes. In addition to aphids [58] and ants [59], *Geobacillus* spp. sequences have been reported from sandflies [60], mosquitoes [61] and ticks [62]. In mosquitoes, *Geobacillus* spp. were identified as part of the core microbiome of dissected reproductive tracts [61]; whereas in the tick *Dermacentor occidentalis*, it was associated with a greater abundance of *Francisella* relative to *Rickettsia* [62]. These findings indicate that although the genus *Geobacillus* is assumed to be exclusively thermophilic, it may have a potential biological role in disease vectors, suggesting that some strains may actually be mesophilic.

The high prevalence of *Geobacillus* sequences in our individual, but not pooled, chigger data raised important questions about both reagent contamination with bacterial DNA and amplification biases caused by variation in GC content. Several recent studies have highlighted the pitfalls of microbiome studies on low biomass samples, where bacterial DNA present in molecular biology reagents competes very effectively as PCR template with bacterial DNA from the sample itself [63, 64]. Since DNA from more than 180 environmental bacterial genera has been detected in commercially available DNA extraction kits [64], assessing the true impact of laboratory contamination on the ensuing data is extremely challenging. The conservative approach that we used here for the individual chiggers was effective, but led to exclusion of more than half of these samples from downstream analyses.

Pooling appears to be an obvious solution to the problem of low biomass samples [65], but is not without its own drawbacks. Genomic GC content is now well known as a source of bias in 16S rRNA datasets, with higher GC content leading to underrepresentation [66], as we observed here with *Geobacillus* spp. (relatively high median GC of ~52%) in pooled samples. At lower template concentrations, denaturation of DNA appears to have been more efficient, revealing OTUs that would have remained hidden had we only sequenced pools.

## Conclusion

This study emphasises that among human disease vectors, chiggers exhibit some of the most complex ecological relationships [67], with high species diversity and low host specificity contributing to elevated rates of coinfection on individual mammalian hosts. The diverse microbiomes of chiggers add a further layer to the network of potential interactions that *Orientia* is exposed to, and future studies should determine whether some of these commensal bacteria affect chigger vector competence. Moreover, the positive correlation we identified here between chigger species richness and scrub typhus incidence deserves further investigation in other endemic countries, especially in relation to the epidemiology of *Orientia* strain diversity [48].

## Supporting information

Supplemental figures, tables, methods and references

16S amplicon OTU tables

## Acknowledgments

The lead author (KC) was supported by the Mahidol-Liverpool Chamlong Harinasuta PhD Scholarship scheme. Field studies were funded by French Agence Nationale de la Recherche grants ANR-07-BDIV-012 (“CERoPath”) and ANR-11-CPEL-002 (“BiodivHealthSEA”) projects to SM, who was also supported by the award “FutureHealthSEA” (ANR-17-CE35-0003). We thank Sabine Dittrich (Lao-Oxford-Mahosot Hospital Wellcome Trust Research Unit) for supplying soil samples from Laos.

