## Supplemental figures, tables, methods and references for "Ecological and microbiological diversity of chigger mites, including vectors of scrub typhus, on small mammals across stratified habitats in Thailand"

### Table of contents


**Figure S1**. Chigger species accumulation curve of an overall 1,395 examined small mammal hosts…………………………………………………………………………………………………………………… p. 2

**Figure S2**. Bar chart showing positive correlation between chigger species richness in the 13 studied sites and latitudinal gradients in Thailand……………………………………………………………….. p. 2

**Figure S3**. Model-averaged importance terms of independent variables used to explain: (A) chigger species richness and (B) scrub typhus human cases..……………………………………………….. p. 3

**Figure S4.** Analysis of differences in mean 16S rRNA gene copy number of Firmicutes among different sample groups as determined by qPCR ……………………………………………………………….. p. 3

**Figure S5**. Maximum likelihood tree of partial Firmicutes 16S rRNA sequences among different sample groups ……………………………………………………………………………………………………. p. 4

**Figure S6**. Boxplot shows a significant difference (two sample *t*-test) in mean GC content of 16S rRNA gene sequences between individual and pooled chigger samples ………………………….. p. 5

**Figure S7**. Principal coordinates analysis plot created using weighted UniFrac metric…………………………………………………………………………………………………………………………… p. 5

**Table S1**. Infestation status for chiggers on small mammals in 13 studied locations in Thailand during 2008 - 2015..……………………………………………………………………………………….... p. 6

**Table S2**. Infestation status and α-diversity of chiggers on small mammal species……………… p. 7

**Table S3**. The prevalence (%) and infestation details of 38 trombiculid species found on small mammals (18 species) in Thailand ………………………………………………………………………………… p. 8

**Table S4**. Bipartite network parameters of host-chigger interaction in the 13 studied sites in Thailand……………………………………………………………………………………………………………………….. p. 9

**Table S5**. Comparison of the generalized linear models (GLM) testing the effect of various independent variables on individual chigger species richness (GLM with Poisson distribution) p. 10

**Table S6**. Generalized linear models of individual chigger species richness (with Poisson distribution)……………………………………………………………………………………………………………………… p. 11

**Table S7**. Human scrub typhus case number and environmental information at the district level for 13 studied sites in Thailand………………………………………………………………………………… p. 12

**Table S8**. Comparison of the general linear models (GLM) to test the effect of various independent variables to scrub typhus human case number (GLM with Poisson distribution). p. 13

**Table S9**. The 34 dominant OTUs (more than 5% proportional read count in a single control) present in background controls……………………………………………………………………………………… p. 14

**Table S10**. α-diversity estimation of bacterial OTUs among sample groups and categories…… p. 15

**Supplemental Materials and Methods** p. 16

**References** p. 23

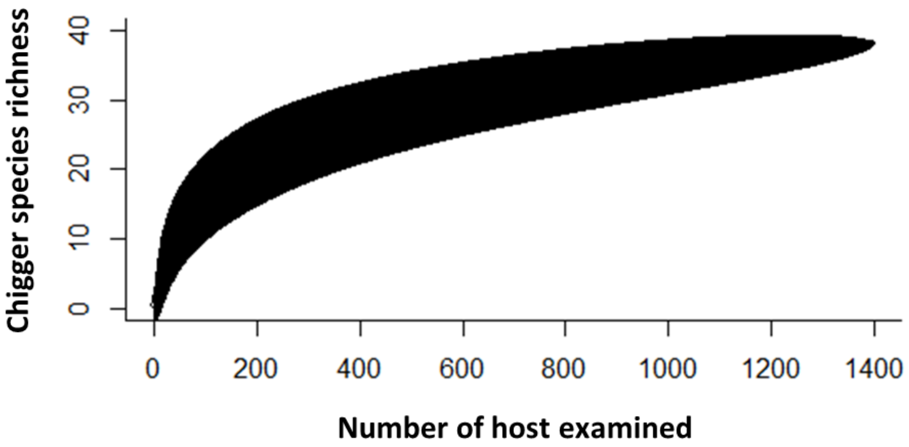


**Figure S1** Chigger species accumulation curve of an overall 1,395 examined small mammal hosts.

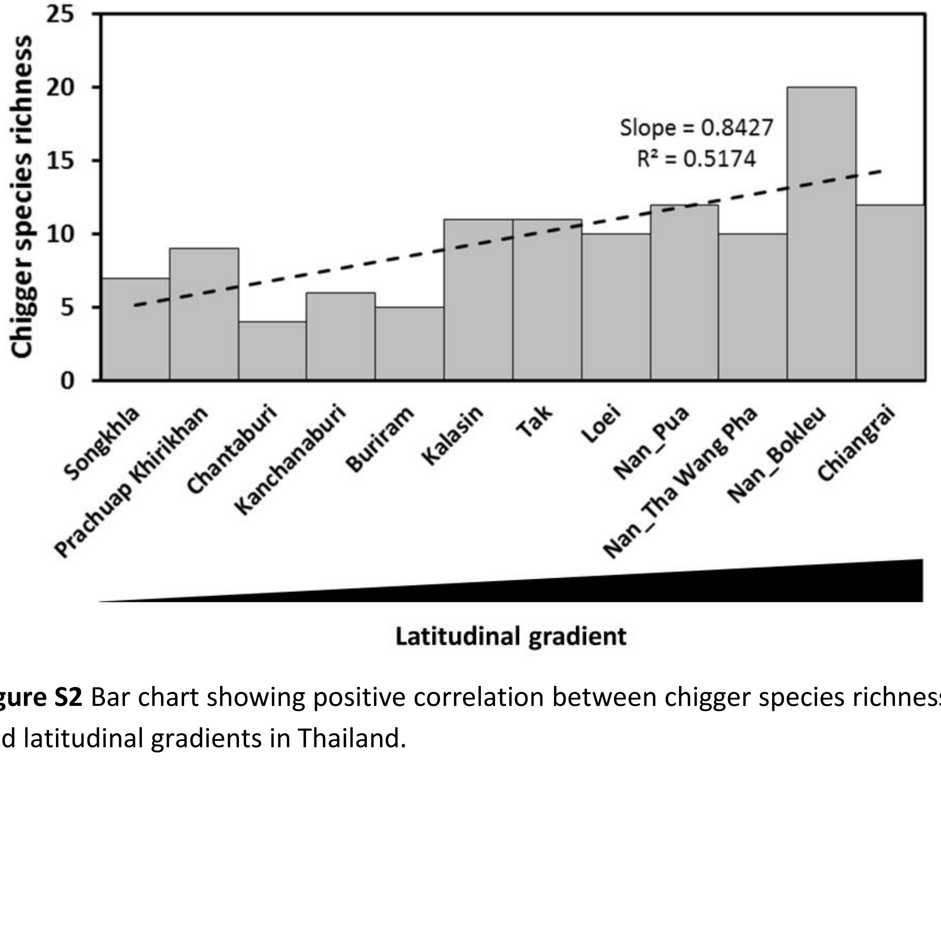


**Figure S2** Bar chart showing positive correlation between chigger species richness in the 13 studied sites and latitudinal gradients in Thailand.

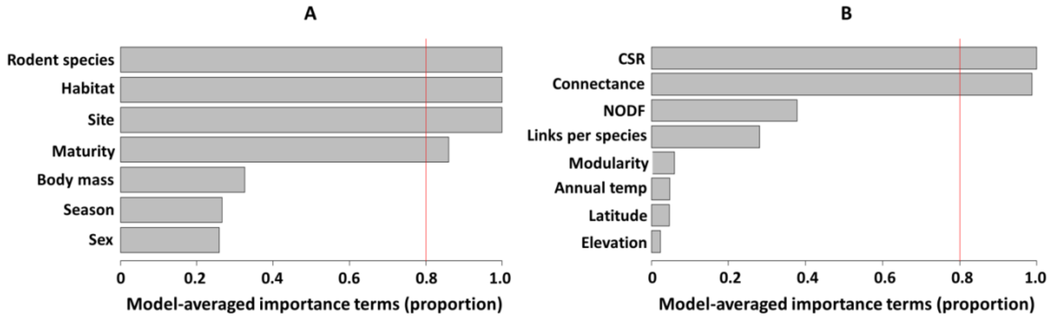


**Figure S3** Model-averaged importance terms of independent variables used to explain: (A) chigger species richness and (B) scrub typhus human cases. The variables with an importance score of >80% proportion support (default in the "gmulti" package in R) were included in the final model.

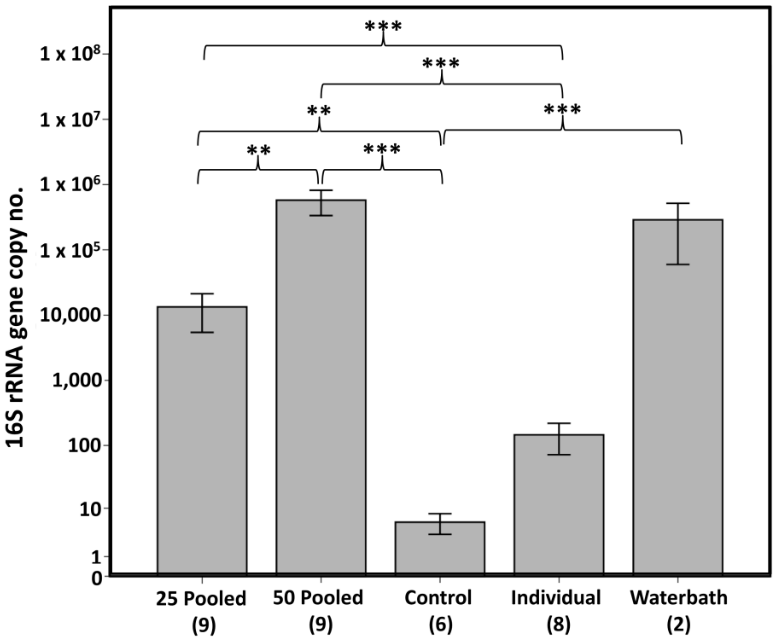


**Figure S4** Analysis of differences in mean 16S rRNA gene copy number of Firmicutes among different sample groups as determined by qPCR (multiple pairwise comparisons after ANOVA with Tukey HSD correction post-hoc test). Numbers in brackets indicate the sample size of each group. (**) $P < 0.01$, (***) $P < 0.001$.

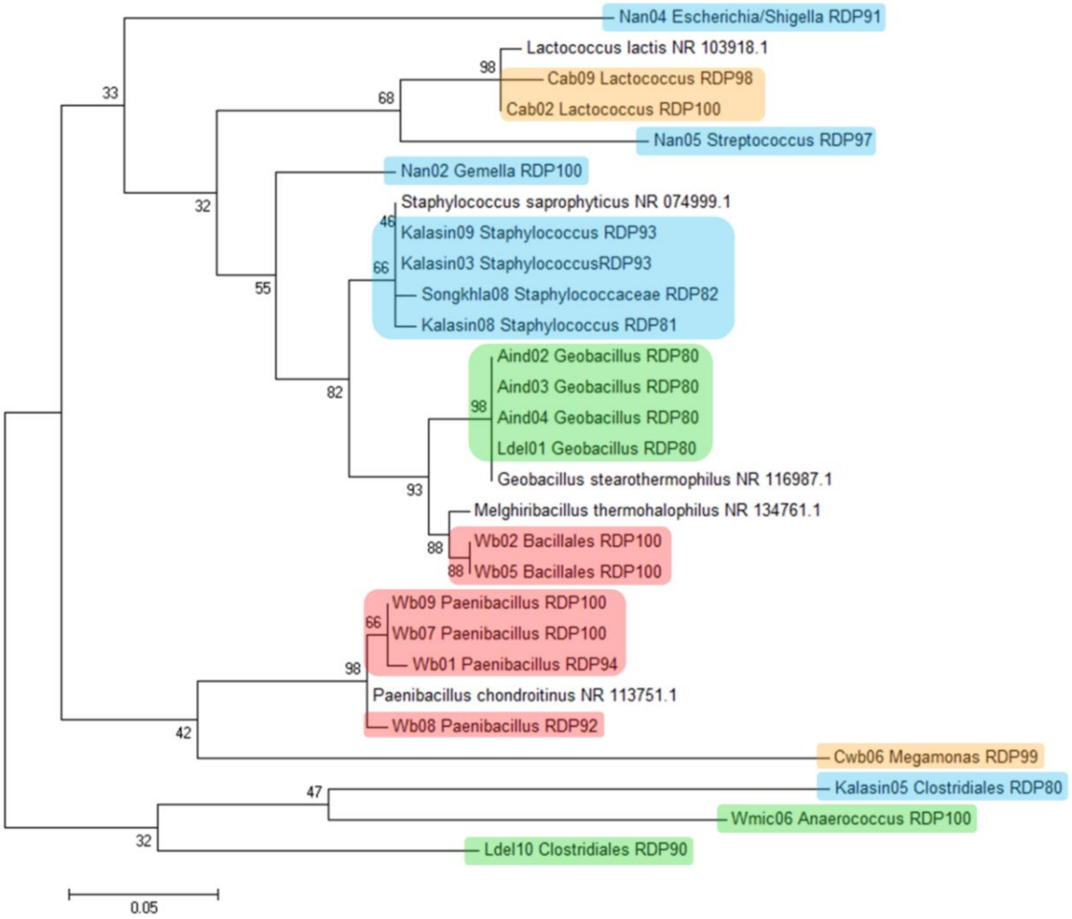


**Figure S5**. Maximum likelihood tree of partial Firmicutes 16S rRNA sequences among different sample groups (red, water bath samples; green, individual chiggers; blue, pooled chiggers; orange; negative controls). Bacterial taxonomic assignment was given in each sequence with a confidence threshold of >80%. Sequences of *Geobacillus stearothermophilus* (accession NR116987.1), *Lactococcus lactis* (NR103918.1), *Melghiribacillus thermohalophilus* (NR134761.1), *Paenibacillus chondroitinus* (NR113751.1) and *Staphylococcus saprophyticus* (NR074999.1) obtained from the National Center for Biotechnology Information were included for comparison. Bootstrap values based on 1,000 replicates are presented at the nodes. The scale bar measures evolutionary distance indicating substitutions per nucleotide.

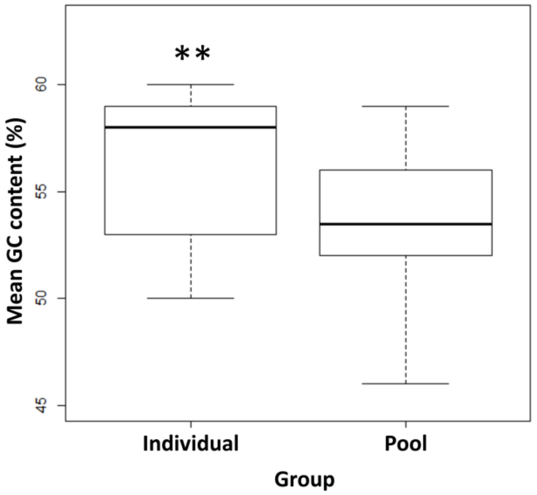


**Figure S6**. Boxplot shows a significant difference (two sample *t*-test) in mean GC content of 16S rRNA gene sequences between individual and pooled chigger samples. (**) $P < 0.01$.

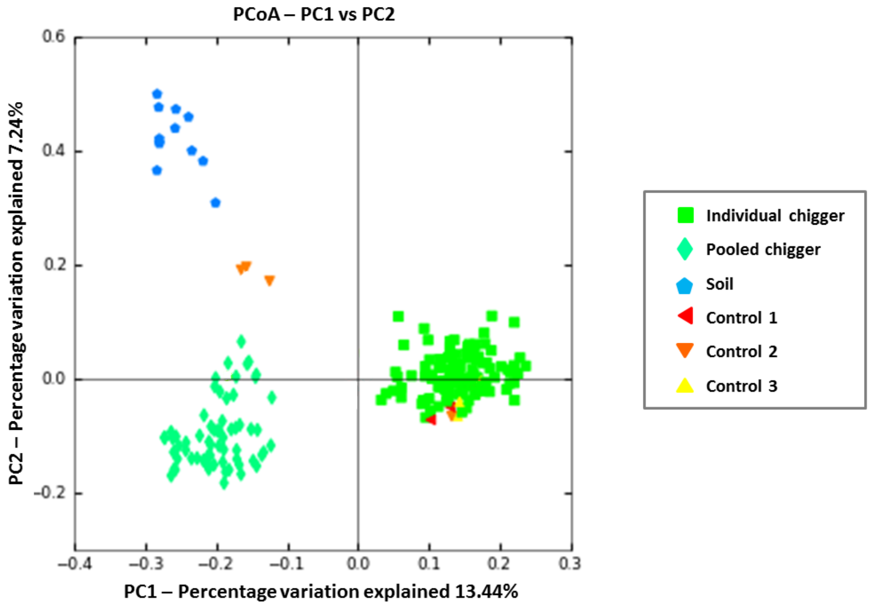


**Figure S7**. Principal coordinates analysis plot created using weighted UniFrac metric. Separation among sample groups is significant (ANOSIM: $R = 0.7997$, $P = 0.001$). Control 1: water in contact with equipment (*i.e.,* glass slides, coverslips, paintbrushes, dissecting needles, gloves and working area) followed by DNA extraction; control 2: nuclease-free water followed by DNA extraction; control 3: nuclease-free water without DNA extraction.

**Table S1** Infestation status for chiggers on small mammals in 13 studied locations in Thailand during 2008 - 2015.

| Province | District | Year | Season | Latitude | Longitude | Number examined animals | Number infested animals | Infestation rate (%) |
|---|---|---|---|---|---|---|---|---|
| Nan | Pua | 2008, 2010 | Dry & Wet | 19.12545 | 100.86202 | 298 | 32 | 10.7 |
| Buriram | Muang | 2009 | Wet | 14.90311 | 103.11365 | 110 | 25 | 22.7 |
| Loei | Muang | 2009 | Wet | 17.45114 | 101.64634 | 232 | 46 | 19.8 |
| Kalasin | Sahatsakhan | 2010 | Dry | 16.29887 | 103.55315 | 186 | 29 | 15.6 |
| Kanchanaburi | Sai Yok | 2011 | Wet | 14.01667 | 99.53333 | 226 | 13 | 5.7 |
| Chiangrai | Wiang Chai | 2011 | Dry | 19.88956 | 99.95113 | 71 | 20 | 28.2 |
| Prachuap Khirikhan | Muang | 2012 | Dry | 11.76527 | 99.65642 | 130 | 39 | 30.0 |
| Nakhonsawan | Tak Fah | 2013 | Wet | 15.34976 | 100.49193 | 87 | 42 | 48.2 |
| Songkhla | Hat Yai | 2013 | Wet | 7.00201 | 100.52691 | 76 | 49 | 64.5 |
| Tak | Mae Sot | 2013 | Dry | 16.80552 | 98.74550 | 37 | 16 | 43.2 |
| Nan | Tha Wang Pha | 2013 | Dry | 19.13926 | 100.71925 | 25 | 10 | 40.0 |
| Nan | Bo Klua | 2014 | Dry | 19.14333 | 101.15395 | 20 | 19 | 95.0 |
| Chantaburi | Laem Singh | 2015 | Wet | 12.50766 | 102.13257 | 76 | 35 | 46.1 |
| | | **Total** | | | | **1,574** | **375** | **23.8** |

**Table S2** Infestation status and α-diversity (CSR = Chigger Species Richness, *H'* = Shannon's index) of chiggers on small mammal species.

| Small mammal host | Number infested hosts | Chigger intensity | Mean intensity | Range | CSR | *H'* |
|---|---|---|---|---|---|---|
| *Bandicota indica* | 87 | 3,297 | 37.9 | 2-238 | 21 | 2.62 |
| *Bandicota savilei* | 3 | 180 | 60.0 | 32-109 | 6 | 1.74 |
| *Berylmys berdmorei* | 6 | 141 | 23.5 | 3-76 | 7 | 1.83 |
| *Berylmys bowersi* | 3 | 340 | 113.3 | 11-317 | 4 | 0.69 |
| *Hylomys suilus* | 3 | 49 | 16.3 | 8-32 | 5 | 1.56 |
| *Leopoldamys edwardsi* | 1 | 12 | 12.0 | 12 | 1 | 0 |
| *Leopoldamys sabanus* | 2 | 30 | 15.0 | 3-27 | 2 | 0.63 |
| *Maxomys surifer* | 20 | 615 | 30.6 | 3-82 | 9 | 1.69 |
| *Mus caroli* | 8 | 422 | 52.6 | 15-156 | 2 | 0.58 |
| *Mus cervicolor* | 14 | 703 | 50.2 | 7-266 | 3 | 0.67 |
| *Mus cookie* | 10 | 292 | 29.2 | 3-73 | 4 | 1.26 |
| *Mus* sp. | 2 | 120 | 60.0 | 24-96 | 1 | 0 |
| *Niviventer fulvescens* | 7 | 103 | 14.7 | 2-34 | 4 | 1.31 |
| *Rattus andamanensis* | 7 | 439 | 62.7 | 8-141 | 12 | 2.26 |
| *Rattus argentiventer* | 6 | 165 | 27.5 | 10-46 | 4 | 1.33 |
| *Rattus exulans* | 5 | 83 | 16.6 | 2-26 | 4 | 1.28 |
| *Rattus sakaeratensis* | 25 | 921 | 36.8 | 4-115 | 10 | 1.87 |
| *Rattus* sp. | 2 | 90 | 45.0 | 2-88 | 1 | 0 |
| *Rattus tanezumi* | 161 | 8,496 | 52.7 | 1-645 | 21 | 2.34 |
| *Tupaia glis* | 3 | 263 | 87.6 | 20-135 | 11 | 2.29 |

**Table S3** The prevalence (%) and infestation details of 38 trombiculid species found on small mammals (18 species) in Thailand.

| Chigger species | Number host species infested | Number host individuals infested | Prevalence (%) |
|---|---|---|---|
| **Tribe Gahrliepiini** | | | |
| *Gahrliepia elbeli* | 3 | 6 | 0.39 |
| *Gahrliepia fletcheri* | 1 | 4 | 0.26 |
| *Gahrliepia* sp., cf. *orientalis* | 2 | 2 | 0.13 |
| *Gahrliepia xiaowoi* | 2 | 3 | 0.20 |
| *Schoengastiella ligula* | 3 | 18 | 1.17 |
| *Walchia chavali* | 3 | 9 | 0.59 |
| *Walchia dismina* | 7 | 13 | 0.85 |
| *Walchia kritochaeta* | 11 | 45 | 2.94 |
| *Walchia lupella* | 5 | 49 | 3.20 |
| *Walchia micropelta* | 12 | 79 | 5.16 |
| *Walchia minuscuta* | 12 | 48 | 3.13 |
| *Walchia pingue* | 7 | 59 | 3.85 |
| *Walchia rustica* | 6 | 23 | 1.50 |
| *Walchia ventralis* | 1 | 1 | 0.07 |
| **Tribe Shoengastiini** | | | |
| *Ascoschoengastia indica* | 8 | 112 | 7.31 |
| *Helenicula kohlsi* | 5 | 12 | 0.78 |
| *Helenicula naresuani* | 2 | 2 | 0.13 |
| *Helenicula pilosa* | 3 | 11 | 0.72 |
| *Helenicula simena* | 3 | 8 | 0.52 |
| *Schoengastia propria* | 1 | 3 | 0.20 |
| *Schoutedenihia centralkwangtunga* | 1 | 2 | 0.13 |
| **Tribe Trombiculini** | | | |
| *Blankaartia acuscutellaris* | 3 | 24 | 1.57 |
| *Leptotrombidium deliense* | 10 | 80 | 5.22 |
| *Leptotrombidium elisbergi* | 1 | 1 | 0.07 |
| *Leptotrombidium imphalum* | 4 | 8 | 0.52 |
| *Leptotrombidium macacum* | 1 | 1 | 0.07 |
| *Leptotrombidium sialkotense* | 1 | 1 | 0.07 |
| *Leptotrombidium* sp., cf. *guzhangense* | 1 | 5 | 0.33 |
| *Leptotrombidium* sp., cf. *maccacum* | 1 | 1 | 0.07 |
| *Leptotrombidium subangulare* | 1 | 1 | 0.07 |
| *Leptotrombidium tenompaki* | 2 | 6 | 0.39 |
| *Leptotrombidium turdicola* | 3 | 7 | 0.46 |
| *Leptotrombidium yunlingense* | 1 | 1 | 0.07 |
| *Lorillatum hekouensis* | 1 | 1 | 0.07 |
| *Microtrombiula munda* | 1 | 3 | 0.20 |
| *Trombiculindus kosapani* n. sp. | 1 | 2 | 0.13 |
| *Trombiculindus paniculatum* | 1 | 1 | 0.07 |
| *Trombiculindus variaculum* | 1 | 1 | 0.07 |

**Table S4** Bipartite network parameters of host-chigger interaction in the 13 studied sites in Thailand (CSR, chigger species richness; NODF, nestedness metric based on overlap and decreasing fill).

| Location | No. hosts examined | No. hosts infested | CSR | NODF | Connectance | Links per species | Modularity |
|---|---|---|---|---|---|---|---|
| Buriram | 131 | 25 | 5 | 32.97 | 0.296 | 1.233 | 0.468 |
| Chantaburi | 76 | 35 | 4 | 15.16 | 0.271 | 0.974 | 0.263 |
| Chiangrai | 70 | 20 | 12 | 14.97 | 0.129 | 0.968 | 0.646 |
| Kalasin | 185 | 29 | 11 | 30.61 | 0.175 | 1.4 | 0.456 |
| Kanchanaburi | 214 | 13 | 6 | 16.66 | 0.231 | 0.947 | 0.617 |
| Loei | 206 | 46 | 10 | 32.81 | 0.176 | 1.446 | 0.372 |
| Nakhonsawan | 87 | 42 | 4 | 56.18 | 0.416 | 1.521 | 0.319 |
| Nan (Bokleu) | 20 | 19 | 20 | 22.36 | 0.131 | 1.282 | 0.536 |
| Nan (Pua) | 138 | 32 | 12 | 22.09 | 0.158 | 1.386 | 0.492 |
| Nan (Tha Wang Pha) | 25 | 10 | 10 | 25.1 | 0.21 | 1.05 | 0.517 |
| Prachuap Khirikhan | 130 | 39 | 9 | 44.82 | 0.199 | 1.458 | 0.249 |
| Songkhla | 76 | 49 | 7 | 40.74 | 0.259 | 1.589 | 0.293 |
| Tak | 37 | 16 | 11 | 24.9 | 0.176 | 1.148 | 0.553 |

**Table S5** Comparison of the generalized linear models (GLM) testing the effect of various independent variables on individual chigger species richness (GLM with Poisson distribution). Selection of the models was done using Akaike's Information Criterion corrected for sample size (AICc). Only the first 10 models are showed. The initial model for AICc selection was Chigger species richness ~ Rodent Species + Sex + Maturity + Weight + Site + Habitat + Season. *K* = the number of estimated variables; log-like, maximized value of the logarithm of the likelihood function; ΔAICc = the difference between AICc value of a given model and the model with minimum AICc; and $W_r$, Akaike weights. Analysis of deviance (ANOVA type II test), significance level (* <0.05, ** < 0.01, *** <0.001). The overall best-supported model is highlighted in bold.

| Model | Dependent variable ~ Independent variables | *K* | Log-like | AICc | ΔAICc | $W_r$ |
|---|---|---|---|---|---|---|
| **1** | **Chigger species richness ~ Rodent Species*** + Maturity* + Site*** + Habitat***** | **5** | **-1302.503** | **2683.754** | **0** | **0.32932** |
| 2 | Chigger species richness ~ Rodent Species*** + Maturity* + Site*** + Habitat*** + Body mass | 6 | -1302.291 | 2685.476 | 1.722 | 0.13921 |
| 3 | Chigger species richness ~ Rodent Species*** + Maturity* + Site*** + Habitat*** + Season | 6 | -1302.446 | 2685.786 | 2.032 | 0.11920 |
| 4 | Chigger species richness ~ Rodent Species*** + Sex + Maturity* + Site*** + Habitat*** | 6 | -1302.483 | 2685.86 | 2.106 | 0.11487 |
| 5 | Chigger species richness ~ Rodent Species*** + Maturity* + Site*** + Habitat*** + Season + Body mass | 7 | -1302.228 | 2687.502 | 3.748 | 0.05054 |
| 6 | Chigger species richness ~ Rodent Species*** + Sex + Maturity* + Site*** + Habitat*** + Body mass | 7 | -1302.271 | 2687.588 | 3.834 | 0.04842 |
| 7 | Chigger species richness ~ Rodent Species*** + Sex + Maturity* + Site*** + Habitat*** + Season | 7 | -1302.425 | 2687.896 | 4.142 | 0.04150 |
| 8 | Chigger species richness ~ Rodent Species*** + Site*** + Habitat*** + Body mass | 5 | -1304.674 | 2688.094 | 4.34 | 0.03759 |
| 9 | Chigger species richness ~ Rodent Species*** + Site*** + Habitat*** | 4 | -1305.758 | 2688.119 | 4.365 | 0.03712 |
| 10 | Chigger species richness ~ Rodent Species*** + Sex + Maturity* + Site*** + Habitat*** + Season + Body mass | 8 | -1302.208 | 2689.618 | 5.864 | 0.01755 |

**Table S6.** Generalized linear models of individual chigger species richness (with Poisson distribution). The best-fit models were selected, and an analysis of deviance table (Type II tests) on the explicative independent variables was generated. Only significant values are included. VIF, variance inflation factor.

| Dependent variable | Explicative variables | Category | Log ratio Chi-square (df, P-value) | Estimate | Std. Error | P-value | VIF |
|---|---|---|---|---|---|---|---|
| **Individual chigger species richness** | **Rodent Species** | | **273.900 (21, < 0.0001)** | | | | 1.051 |
| | *B. indica vs* | *B. savilei* | | -0.916 | 0.46 | 0.046 | |
| | | *B. bowersi* | | -1.825 | 0.73 | 0.013 | |
| | | *M. caroli* | | -1.726 | 0.34 | < 0.0001 | |
| | | *M. cervicolor* | | -2.159 | 0.27 | < 0.0001 | |
| | | *M. cookii* | | -1.445 | 0.34 | < 0.0001 | |
| | | *N. fulvescens* | | -1.172 | 0.42 | 0.005 | |
| | | *R. exulans* | | -2.509 | 0.42 | < 0.0001 | |
| | | *T. glis* | | 1.069 | 0.43 | 0.014 | |
| | **Maturity** | | **8.179 (1, 0.004)** | | | | 1.052 |
| | Adult *vs* | Juvenile | | -0.283 | 0.10 | 0.004 | |
| | **Site** | | **50.443 (11, < 0.0001)** | | | | 1.203 |
| | Buriram *vs* | Chantaburi | | -0.702 | 0.32 | 0.033 | |
| | | Nan (Bo Kleu) | | 1.006 | 0.39 | 0.010 | |
| | | Nan (Tha Wang Pha) | | 0.711 | 0.31 | 0.023 | |
| | | Songkhla | | 0.456 | 0.21 | 0.035 | |
| | **Habitat** | | **55.420 (3, < 0.0001)** | | | | 1.159 |
| | Forest *vs* | Settlement | | -1.074 | 0.17 | < 0.0001 | |

**Table S7.** Human scrub typhus case number and environmental information at the district level for 13 studied sites in Thailand.

| Location | District | Latitude | Elevation (m) | Annual mean temp (ºc) | Scrub typhus case number (year) |
|---|---|---|---|---|---|
| Buriram | Muang | 14.90311 | 141.9 | 25 | 1 (2009) |
| Chantaburi | Laem Singh | 12.50766 | 25.73 | 26 | 1 (2015) |
| Chiangrai | Wiang Chai | 19.88956 | 111.7 | 30 | 8 (2011) |
| Kalasin | Sahatsakhan | 16.29887 | 279.5 | 27 | 21 (2010) |
| Kanchanaburi | Sai Yok | 14.01667 | 279.5 | 26 | 1 (2011) |
| Loei | Muang | 17.45114 | 56.26 | 28 | 7 (2009) |
| Nakornsawan | Tak Fah | 15.34976 | Na | Na | 2 (2013) |
| Nan | Bo Kleu | 19.14333 | 447.3 | 20 | 209 (2014) |
| Nan | Pua | 19.12545 | 144 | 26 | 20 (2010) |
| Nan | Tha Wang Pha | 19.13926 | 176.3 | 27 | 37 (2013) |
| Prachuap Khirikhan | Muang | 11.76527 | 111.7 | 26 | 4 (2012) |
| Songkhla | Hat Yai | 7.00201 | 62.9 | 28 | 7 (2013) |
| Tak | Mae Sot | 16.80552 | 371.6 | 23 | 20 (2013) |

**Table S8.** Comparison of the general linear models (GLM) to test the effect of various independent variables to scrub typhus human case number (GLM with Poisson distribution). Selection of the models was done using Akaike's Information Criterion corrected for sample size (AICc). Only the top 10 models are shown. The initial model for AICc selection was Scrub Typhus Human Case Number ~ CSR + Latitude + Elevation + Annual Mean Temp + Network Modularity + NODF + Links per species + Network Connectance. *K*, the number of estimated variables; log-like, maximized value of the logarithm of the likelihood function; ΔAICc, the difference between AICc value of a given model and the model with minimum AICc; $W_r$, Akaike weights; CSR = chigger species richness; NODF = nestedness. Analysis of deviance (ANOVA type II test) significance level (* <0.05, ** < 0.01, *** <0.001). The overall best-supported model is highlighted in bold.

| Model | Dependent variable ~ Independent variables | *K* | Log-like | AICc | ∆AICc | $W_r$ |
|---|---|---|---|---|---|---|
| 1 | Scrub typhus human case number ~ CSR*** + NODF* + Network Connectance*** | 4 | -47.289 | 114.578 | 0 | 0.3244 |
| **2** | **Scrub typhus human case number ~ CSR*** + Network Connectance*** ** | **3** | **-50.626** | **114.966** | **0.388** | **0.2672** |
| 3 | Scrub typhus human case number ~ CSR*** + Links per species* + Network Connectance*** | 4 | -47.639 | 115.279 | 0.701 | 0.2285 |
| 4 | Scrub typhus human case number ~ CSR*** + Annual Mean Temperature +Network Connectance** | 4 | -49.775 | 119.55 | 4.972 | 0.027 |
| 5 | Scrub typhus human case number ~ CSR*** + Network Modularity + Links per species** + Network Connectance*** | 5 | -45.43 | 119.661 | 5.083 | 0.0255 |
| 6 | Scrub typhus human case number ~ CSR*** + Network Modularity + NODF** + Network Connectance*** | 5 | -45.679 | 120.159 | 5.581 | 0.0199 |
| 7 | Scrub typhus human case number ~ CSR*** + Latitude + NODF** + Network Connectance*** | 5 | -45.78 | 120.36 | 5.782 | 0.0181 |
| 8 | Scrub typhus human case number ~ CSR*** + Elevation + Network Connectance*** | 4 | -50.349 | 120.699 | 6.121 | 0.0152 |
| 9 | Scrub typhus human case number ~ CSR*** + Latitude + Links per species** +Network Connectance*** | 5 | -46.006 | 120.12 | 5.542 | 0.0143 |
| 10 | Scrub typhus human case number ~ CSR*** + Network Modularity + Netwrok Connectance*** | 4 | -50.565 | 121.13 | 6.552 | 0.0122 |

**Table S9.** The 34 dominant OTUs (more than 5% proportional read count in a single control) present in background controls. Control 1: water in contact with equipment (i.e., glass slides, coverslips, paintbrushes, dissecting needles, gloves and working area) followed by DNA extraction; control 2: nuclease-free water followed by DNA extraction; control 3: nuclease-free water without DNA extraction. The highest OTU proportions in particular controls are showed in bold. The following OTUs were previously reported as potential contaminants in Tanner et al. [1][a], Grahn et al. [2][b], Barton et al. [3][c], Salter et al. [4][d], Galan et al. [5][e], and Glassing et al. [6][f].

| Bacterial taxa (OTU identifier) | Maximum proportion (%) | | | |
|---|---|---|---|---|
| | All controls | Control 1 | Control 2 | Control 3 |
| Species *Luteibacter rhizovicinus* (177555) | 41.59 | 24.73 | **41.59** | 0.07 |
| Genus *Flavobacterium* (4438548)[d] | 22.86 | 0.00 | **22.86** | 0.03 |
| Genus *Fimbriimonas* (New.ReferenceOTU3) | 20.36 | 0.00 | 3.54 | **20.36** |
| Family Oxalobacteraceae (4476547)[f] | 19.15 | 4.64 | 12.50 | **19.15** |
| Genus *Stenotrophomonas* (2806353)[a,b,d,f] | 18.02 | 14.22 | 3.30 | **18.02** |
| Species *Bacillus thermoamylovorans* (21214) | 14.97 | - | **14.97** | - |
| Family Comamonadaceae (4396454)[f] | 14.92 | 3.31 | 9.08 | **14.92** |
| Genus *Sphingomonas* (4423410)[a,d] | 14.59 | 8.72 | 2.47 | **14.59** |
| Family Intrasporangiaceae (4432889) | 13.81 | 7.95 | 1.22 | **13.81** |
| Genus *Streptococcus* (4473883)[d,e,f] | 12.89 | - | 0.56 | **12.89** |
| Genus *Hydrogenophilus* (4317875) | 12.85 | - | **12.85** | 0.03 |
| Genus *Pseudomonas* (4456891)[b,d,e,f] | 12.49 | 11.64 | 5.59 | **12.49** |
| Genus *Novosphingobium* (581019)[d] | 12.28 | 5.70 | 4.21 | **12.8** |
| Family Cytophagaceae (737260) | 10.67 | - | **10.67** | 0.02 |
| Species *Methylobacterium organophilum* (New.ReferenceOTU2) | 10.56 | **10.56** | 10.15 | 0.03 |
| Family Caulobacteraceae (4353264) | 10.41 | 6.76 | **10.41** | 4.80 |
| Phylum Cyanobacteria (98258) | 10.06 | - | 0.04 | **10.06** |
| Genus *Ochrobactrum* (4388385)[d] | 9.84 | **9.84** | 1.25 | 9.53 |
| Genus *Rhodococcus* (4468125)[d] | 9.44 | 3.42 | 7.32 | **9.44** |
| Genus *Wautersiella* (New.ReferenceOTU2903)[d] | 8.92 | - | **8.92** | - |
| Genus *Cloacibacterium* (4154872)[f] | 8.64 | - | **8.64** | - |
| Family Methylophilaceae (101445) | 7.87 | 1.85 | 0.41 | **7.87** |
| Genus *Lactobacillus* (619224)[f] | 7.32 | - | - | **7.32** |
| Family Caulobacteraceae (4339358) | 7.23 | 6.74 | **7.23** | 0.01 |
| Genus *Corynebacterium* (942245)[d,e,f] | 7.13 | - | **7.13** | - |
| Genus *Acinetobacter* (4482374)[a,c,d,e] | 7.11 | 0.03 | **7.11** | 3.56 |
| Genus *Fluviicola* (646052) | 6.91 | - | **6.91** | 0.03 |
| Class Alphaproteobacteria (New.CleanUp.ReferenceOTU210) | 6.07 | - | **6.07** | 0.03 |
| Family Microbacteriaceae (1109043) | 6.06 | - | **6.06** | - |
| Family Caulobacteraceae (810889) | 5.84 | - | **5.84** | 0.61 |
| Species *Kocuria rhizophila* (4477552) | 5.49 | 3.35 | **5.49** | 2.00 |
| Family Bradyrhizobiaceae (4475561) | 5.14 | - | 0.45 | **5.14** |
| Family Polyangiaceae (New.ReferenceOTU9) | 5.03 | - | 0.03 | **5.03** |

**Table S10.** α-diversity estimation of bacterial OTUs among sample groups and categories.

| Sample group (*n*) | Sample categories (*n*) | Observed (SD) | Chao1 (SD) | PD_whole_tree (SD) |
|---|---|---|---|---|
| Individual chigger (109) | | 79.6 (14.7) | 95.7 (18.7) | 9.3 (1.5) |
| Chigger species: | *A. indica* (7) | 90 (14.7) | 111.1 (12.6) | 10.1 (2.0) |
| | *B. acuscutellaris* (7) | 71.1 (8.9) | 83.7 (12.5) | 8.2 (1.1) |
| | *H. kohlsi* (11) | 73.5 (12.8) | 89.4 (20.5) | 9.3 (1.5) |
| | *H. pilosa* (10) | 85.3 (9.6) | 102.0 (15.6) | 9.9 (1.4) |
| | *L. deliense* (39) | 78.0 (16.2) | 94.1 (20.1) | 9.1 (1.4) |
| | *S. ligula* (10) | 78.0 (11.6) | 92.1 (16.7) | 8.4 (0.8) |
| | *W. kritochaeta* (12) | 76.8 (16.4) | 96.2 (23.1) | 8.9 (1.7) |
| | *W. micropelta* (13) | 87.9 (12.2) | 101.6 (11.5) | 10.7 (1.4) |
| Habitat: | Forest (9) | 74.2 (8.1) | 93.8 (14.8) | 8.7 (0.7) |
| | Dry land (48) | 80.3 (15.2) | 96.6 (19.2) | 9.5 (1.5) |
| | Rain-fed land (28) | 78.4 (11.9) | 93.7 (17.1) | 9.0 (1.4) |
| | Settlement (18) | 79.9 (17.6) | 96.0 (21.6) | 9.3 (1.7) |
| Site: | Buriram (9) | 89.6 (16.2) | 103.3 (18.7) | 10.5 (1.7) |
| | Chiangrai (7) | 78.1 (13.2) | 91.6 (21.3) | 8.6 (0.9) |
| | Kalasin (12) | 85.1 (14.1) | 103.4 (20.9) | 9.6 (1.2) |
| | Kanchanaburi (2) | 83.0 (1.4) | 90.5 (0.4) | 10.0 (0.1) |
| | Loei (17) | 80.9 (15.4) | 95.7 (19.3) | 9.5 (1.5) |
| | Nakhonsawan (3) | 67.0 (6.2) | 82.2 (5.2) | 7.8 (0.5) |
| | Nan (10) | 81.1 (16.8) | 98.5 (20.5) | 9.2 (2.3) |
| | Prachuab Kirikhan (17) | 73.0 (15.2) | 92.7 (17.1) | 8.9 (1.5) |
| | Songkhla (15) | 73.8 (13.2) | 89.7 (17.1) | 8.6 (1.1) |
| | Tak (17) | 82.2 (12.0) | 97.6 (19.3) | 9.9 (1.4) |
| Pooled chigger (65) | | 108.8 (29.3) | 132.2 (31.6) | 11.3 (2.8) |
| Chigger species: | *A. indica* (8) | 101.7 (33.7) | 128.3 (38.9) | 11.3 (3.8) |
| | *L. deliense* (12) | 87.9 (22.2) | 114.9 (16.2) | 9.7 (1.8) |
| | *W. micropelta* (8) | 124.2 (9.9) | 151.2 (9.4) | 12.7 (1.4) |
| | *W. minuscuta* (2) | 178.5 (48.8) | 195.7 (47.1) | 18.2 (3.4) |
| Habitat: | Forest (3) | 74.0 (13.8) | 104.7 (6.1) | 8.2 (1.3) |
| | Dry land (4) | 130.0 (11.7) | 147.4 (8.9) | 13.2 (1.0) |
| | Rain-fed land (7) | 102.1 (18.1) | 131.0 (19.7) | 11.0 (1.6) |
| | Settlement (4) | 121.7 (29.5) | 146.7 (30.7) | 13.3 (3.2) |
| Site: | Buriram (5) | 73.6 (13.7) | 106.2 (24.5) | 8.2 (1.3) |
| | Chantaburi (3) | 117.0 (12.1) | 168.1 (23.2) | 13.0 (1.5) |
| | Chiangrai (3) | 101.3 (18.1) | 125.3 (23.2) | 10.3 (2.2) |
| | Kalasin (3) | 97.6 (12.0) | 108.4 (24.5) | 9.8 (1.0) |
| | Kanchanaburi (3) | 90.3 (16.0) | 102.8 (20.6) | 9.5 (1.8) |
| | Loei (3) | 113.6 (11.0) | 129.0 (10.8) | 11.1 (0.3) |
| | Nakhonsawan (3) | 110.6 (13.7) | 125.2 (18.2) | 10.6 (1.3) |
| | Nan (3) | 165.6 (11.0) | 182.1 (9.7) | 16.1 (0.8) |
| | Prachuab Kirikhan (3) | 124.0 (13.0) | 149.9 (7.8) | 12.6 (1.3) |
| | Songkhla (3) | 97.3 (8.5) | 123.2 (20.2) | 10.1 (1.1) |
| | Tak (3) | 122.3 (9.5) | 148.9 (16.9) | 12.3 (0.9) |
| Soil (12) | | 109.8 (34.3) | 150.6 (38.6) | 13.1 (3.0) |
| Background control (15) | | 80.5 (33.7) | 103.4 (40.9) | 10.1 (3.8) |

**Supplemental Materials and Methods**

*Trapping of small mammals and chigger collections*

This study utilised chigger material collected previously for a taxonomic study in Thailand [7]. In brief, small mammals were trapped across 13 localities between 2008 – 2015, once each in the dry season and wet season. The animals were euthanized and identified morphologically using taxonomic keys or by molecular barcoding of the *coi* gene, then weighed and sexed. Maturity of the animals and the sex of juveniles was gauged during dissection by examination of the internal reproductive organs (presence of ovaries, uterus, testes or seminal vesicle). Chiggers were removed from the mammal cadavers and fixed in 70 - 95% ethanol. Mites collected from the same animals were placed in a single tube and counted to estimate the intensity and abundance of infestation, as defined by Rózsa *et al*. [8]. To identify and estimate chigger species richness, 10 - 20% of chiggers from each infested animal were intentionally selected by differences in observed sizes and microscopic appearance as subsamples representative for a specific animal. The mites were cleared in Berlese's fluid and identified morphologically as described previously [7].

*Ecological analysis*

For ecological analysis, trapping sites were divided equally into four different types of habitats with respect to human land use (anthropization index), spanning low to high levels of disturbance [9-11]. These were (a) forest - including primary, secondary or community forest with mature plantations of timber woods (*e.g*., teak, rubber, or eucalyptus); (b) dry land – including non-flooded agricultural land (*e.g*., cassava, maize, pineapple or dry rice fields) and fallow land, grasslands, dry fields and shrub; (c) rain-fed land - flooded farmland and cultivated land (*e.g*., rice fields, legume, shrimp or fish farm) including other types of floodable land, swamp or marsh; (d) settlement - built-up areas (*e.g*., an isolated house, factory, market, village or city).

Mean intensity of conspecific chigger species living on an infested host and the range of chigger infestation on the small mammals were estimated. Chigger species richness (CSR), first-ordered Jack-knife (Jack1) and Shannon index (H') were calculated across different habitats, seasons, study sites and host attributes (sex and maturity) using the "BiodiversityR" package [12] implemented in the R freeware programming environment [13]. Chigger species accumulation curves were generated for evaluation of small mammal sample size adequacy, as well as to illustrate differences in chigger species richness among different factors. Nonparametric Kruskal–Wallis and multiple pairwise comparison tests were performed to investigate the effects of habitat on chigger species richness.

Twelve chigger species were included in an analysis of association with habitat type; these were selected from chigger species that infested ≥10 individual hosts. To visualize the association between the chigger species and habitat types, correspondence analysis (CA) was performed using the "FactoMineR" package in R freeware [14]. This CA is similar to principle component analysis (PCA), a statistical method to simplify the two categorical variables (chigger species and habitats in this case) in the dataset by reducing dimensionality of the dataset and to visualize the association of two particular variables through a two-dimensional plot, whereas PCA is used to analyse numerical variables [14].

*Network analyses of host-chigger interactions*

Host-chigger network analysis was conducted to explore interactions through assessment of network architectures (properties) by focusing at both the level of host species (pooled host species and locations) and the host individual. To study the community ecology of host-chigger interactions, bipartite network analyses were conducted on both the community and individual basis of the host-ectoparasite interactions using “vegan” [15] and “bipartite” packages [16] implemented in R freeware. A host-chigger interaction matrix (presence/absence) of the pooled 13 study sites was created, with host species as rows and chigger species as columns. The matrix was visualized for bipartite network and nestedness patterns using “visweb” and “plotweb” functions, respectively. The chigger species specificity index was also computed by the “specieslevel” function to identify specialists or species with few links (lower sharing of the same chigger species with other hosts) and generalists or species with many links (higher sharing of several chigger species with the other hosts) in the community.

Meanwhile, on the host individual basis, the same type of matrices was generated for each studied site/community. Subsequently, a number of network properties were estimated: (1) Nestedness is the degree of how many interactions realized by specialists belong to subset of those realized by generalists in the particular community. The NODF (Nestedness metric based on overlap and decreasing fill) was computed using “nestednodf” functions. The NODF ranges from 0 - 100, where a value of 100 indicates perfect nestedness and 0 represents an absence of nestedness [17]. (2) Connectance is defined as the proportion of possible links between the realized species [18]; in other words, this can be described as the proportion of established interactions or network complexity in a particular community. (3) Links per species is the mean number of the same chigger species shared (links) per host. (4) Modularity is a measure of community structure/compartmentalization of a particular network. The higher the modularity, the more sub-communities dependently clustered in the network. In other words, sub-communities consist of species with many links among themselves and sparsely interact with species in other sub-communities [19]. Network modularity of the 13 sites was computed by the “computeModules” function, where each matrix of bipartite weighted graph is taken into account to compute the parameter. A repeat of 100 000 computational steps was applied by default to confirm that no better clustering than the current one could be identified [16].

Bipartite networks were transformed to unipartite networks using the “tnet” package [20]. Unipartite network plots illustrate the relative interaction patterns among hosts regarding the co-occurrence of chigger species shared within the particular community. An Eigen value of centrality was calculated using the “evcent” function from the “igraph” package [21]. This centrality measurement was used to estimate the role of each host as a connector to other hosts with respect to the same shared chigger species. A higher value of centrality of a node (host) is associated with a higher connection to the other nodes (other hosts), suggesting a high number of parasite species co-occurred in the network [22].

*Multiple regression models of independent variables explaining chigger species richness*

Generalized linear models (GLM) were constructed in order to identify potential effects of host attributes (species, sex, maturity and body mass) and ecological factors (habitat, site and season) on chigger species richness at the level of individual hosts. Linear regression models using the “Poisson” family for chigger species richness count (skewed data containing many zeros) were modelled in the “lme4” package [23] embedded in R freeware. A selection of models based on the likelihood-based method, Akaike’s Information Criterion (AICc), was adjusted for sample size using the “gmulti” package [24] in R freeware. Model selection can sometimes produce an uncertain fit or implausible model in the output. Accordingly, the quality of various models was quantified by the model weight value (Akaike's weight: Wr), which can be realized as the probability that a particular model is the best selection from the available data [25]. Delta AICc (ΔAICc), the difference between the AICc value of a given model and a model with minimum AICc, was also calculated in order to facilitate the best model selection. Model-averaged Importance of Term (MaIT) was illustrated through histogram plotting to identify the significance of particular variables. The MaIT is defined as the proportion of the best models in which each given candidate variable term appears after all possible models are computed (*i.e.*, 150 models for chigger species richness and 250 models for scrub typhus incidence in the present analysis) [26]. The variables with an importance score of >80% proportion support (default in the "gmulti" package) were included in the final model.

The best-selected model was evaluated from following criteria: low AICc score, clear ΔAICc difference from the substitute models, high Wr score, and explicative variables passing with 80% MaIT support. The best model was subsequently designated for an Analysis of Deviance table (ANOVA type II test) in order to emphasize the role of explicative independent variables on individual chigger species richness. To identify the degree of multicollinearity among explicative variables, the variance inflation factor (VIF) was computed in the "car" package [27] in R freeware. The higher the VIF value, the stronger the collinearity, and a VIF score >10 was used as a common cut-off threshold to indicate strong multicollinearity in a model [28].

Data for scrub typhus human case numbers from the 13 studied sites were obtained from the Bureau of Epidemiology, Ministry of Public Health, Thailand (unpublished data). Cases were recorded in Report 506 (http://www.boe.moph.go.th/boedb/surdata/disease.php?dcontent=def&ds=44) for patients identified as a "probable case" or "confirmed case" following the criteria in the International Classification of Diseases and Related Health Problems [29]. These human case data were selected for the particular districts and years when the field surveys of small mammals had been conducted. Geographical and environmental information; *i.e.*, GPS coordinates (latitude-longitude), elevation, and annual mean temperature of the studied sites were derived from the CERoPath project [30]. The environmental information above and host-chigger network properties were assigned as candidate independent variables to explain scrub typhus human case number across the country.

To examine pairwise relationships between those variables above and scrub typhus case number, the non-parametric Spearman's rank with significance test was conducted in R freeware. Finally, in order to evaluate the best fit model and initially identify constitutive variables accounting for scrub typhus epidemiology in Thailand, GLM with "Poisson" family and model selection with AICc were applied as described above.

*DNA extraction*

As clearing in Berlese's fluid destroys DNA, the identity of chiggers destined for DNA extraction was established using an autofluorescence microscopy method as previously described. Genomic DNA was purified using the DNeasy Blood & Tissue Kit (Qiagen, Hilden, Germany). The mites were crushed with polypropylene pestles in a microcentrifuge tube containing ATL buffer and proteinase K solution (Qiagen). The lysates were incubated at 56°C overnight, and then subsequent steps followed the manufacturer's protocol. A minimal volume (30 µl) of nuclease-free water (Ambion, Thermo Fisher Scientific, Altrincham, UK) was used in the DNA elution step. The DNA concentrations were determined by a double-stranded DNA fluorescence-labelling method (Quant-iT Picogreen;

Invitrogen, Altrincham, UK) read in an Infinite F200 microplate fluorimeter with Magellan data analysis software (Tecan, Männedorf, Switzerland).

*Library preparation and next generation sequencing of 16S rRNA amplicons*

To determine the bacterial microbiome of chiggers, a dual-index nested PCR protocol for MiSeq (Illumina, San Diego, CA, USA) sequencing was applied [31-33] targeting the v4 region of the 16S rRNA gene. The second round PCR to attach the barcode indices and Illumina sequencing adaptors to the first-round amplicon products was performed using the Nextera XT DNA protocol (Illumina). Three types of negative controls were included on every MiSeq run in order to identify potential background contamination from sample manipulation equipment, DNA extraction kits and PCR reagents used in the library preparation steps. These were (a) water in contact with equipment (*i.e.*, glass slides, coverslips, paintbrushes, dissecting needles, gloves and the working area) followed by DNA extraction in the DNeasy Blood & Tissue Kit (Qiagen); (b) nuclease-free water followed by DNA extraction; and (c) nuclease-free water without DNA extraction.

The first round PCR was conducted in a 25 µl reaction containing Applied Biosystems (Warrington, UK) reagents in a T3 thermocycler (Biometra) as follows: 1× GeneAmp PCR buffer I; GeneAmp dNTPs (0.5 mM final concentration); 0.5 µM each 16S v4 forward and reverse primer; 1.25 U AmpliTaq DNA Polymerase Low DNA (LD); and 1 µl of DNA template. The PCR comprised initial denaturation at 94°C for 5 min, followed by 18 cycles of denaturation at 94°C for 1 min, annealing at 56°C for 1 min, extension at 72°C for 1 min, and final extension at 72°C for 5 min. Subsequently, PCR products were size-selected and purified using a Chroma Spin-200 column (Takara-Clontech, Saint-Germain-en-Laye, France) following the manufacturer's protocol. The DNA fragments >150 bp were retained and primer-dimers <50 bp were removed by this step after elution with 50 µl nuclease-free water. The size-selected PCR products were then used as the DNA templates for second-round PCR in a 25 µl reaction containing 1× GeneAmp PCR buffer I; GeneAmp dNTPs (0.5 mM final concentration); 0.5 µM each forward and reverse index barcoding primer; 1.25 U AmpliTaq DNA Polymerase LD; and 3 µl DNA. The PCR comprised initial denaturation at 94°C for 5 min, followed by 25 cycles of denaturation at 94°C for 1 min, annealing at 65°C for 1 min, extension at 72°C for 1 min and final extension at 72°C for 5 min.

The second-round PCR products (5 µl) were visualised on a 1.2% agarose gel incorporating SYBR Safe DNA gel stain (Invitrogen, Altrincham, UK). Gels were visualised using a G:Box Gel Documentation System (Syngene, Cambridge, UK). The PCR products were purified individually using a QIAquick PCR Purification Kit (Qiagen) following the manufacturer's protocol, with 30 µl of nuclease-free water used in the elution step. For each sequencing run, the 96 purified PCR products were pooled by adjusting the DNA volume on the basis of band density in the gel images. A volume of 5, 10 or 15 µl was taken from the samples with high, medium and low band densities, respectively. The concentration of pooled DNA was determined using the Quant-iT Picogreen dsDNA kit (Invitrogen), and then submitted for sequencing with 300 bp paired-end chemistry on the Illumina MiSeq platform at the Centre for Genomic Research (University of Liverpool).

*Quality filtering*

The reads in fastq format were trimmed with CUTADAPT v.1.2.1 [34] and SICKLE v.1.200 [35], and the sequences with an average nucleotide base quality score lower than 20, as well as a length shorter than 10 bp after trimming, were discarded. This produced the forward (R1) and reverse (R2) reads of the read-pairs, whereas the singlet reads (R0; where only member of a read-pair passed filtering) were excluded from further steps. Subsequently, error correction of the reads was performed using the BayesHammer algorithm in SPAdes v.3.1.0 [36, 37]. Read-pairs (R1 and R2) were then aligned using PANDAseq [38], generating assembled reads representative for the certain pairs. Only reads with a size between 270-300 bp were retained. Finally, the aligned reads from the four lanes of MiSeq sequencing were combined in a single fasta file. The file containing the whole dataset from 377 samples was used for further microbiome analyses.

*Microbiome profiling*

Analyses of 16S rRNA microbiome profile were performed using the Quantitative Insights into Microbial Ecology (QIIME) software package, version 1.8.0, http://qiime.org [39]. Operational Taxonomic Units (OTUs) were grouped by sequence similarity or using an open-reference OTU picking approach using the USEARCH61 method [40]. All reads were binned at 97% similarity against the Greengene database v. 13_8 [41]. Any reads that did not match the reference database were subsequently clustered *de novo* against each other with the same similarity threshold. These steps were performed using the "pick_open_reference_otus.py" command, in which bacterial taxonomic assignment with UCLUST against the Greengene database v.13_8 [40], sequence alignment with PyNAST [42], and tree-building with FastTree v.2.1.3 [43] were also generated in the outputs. Within an OTU, the most abundant read was selected as a representative sequence for that particular OTU. The OTU table (providing the taxonomic assignment and read count of 16S rRNA gene sequences in each OTU and sample) was created in biom (Biological Observation Matrix) format.

Chimeric sequences were identified using "ChimeraSlayer" [44] and a new OTU table and phylogenetic tree were generated after their removal. To facilitate a manual investigation of microbiome profiles, the OTU table was filtered to discard any OTUs with a relative abundance (proportion of total read count) of <1% across samples, whereas the OTUs represented by ≥1% of reads were retained for further analyses. The OTUs with fewer than five read counts were identified as singletons and then removed from the OTU table.

There are several potential sources of errors and biases during the sample preparation and generation of a 16S rRNA library; *e.g.*, equipment and reagents used in specimen manipulation and DNA extraction kit and water contamination, which produce substantial challenges for high-throughput sequencing experiments, particularly when dealing with low biomass microbiota samples [4, 6]. As our study dealt with such challenging specimens, very small DNA yields from individual chiggers were obtained, and these low biomass samples could fail to outcompete contaminating 16S rRNA gene sequences. Accordingly, a sample-control similarity check using the β-diversity approach (Bray-Curtis dissimilarity) was applied using the "ecodist" package [45] implemented in R freeware [13]. Any samples that exhibited a microbiome pattern (OTU profile) to the background controls of >20% similarity were excluded, as about a 20 - 30% cut-off seemed acceptable to discriminate samples and controls according to previous studies [46, 47]. We decided to remove low quality samples before conducting comparative analyses rather than subtracting contaminant OTUs of likely background control origin, as the latter could affect the relative abundance in the samples.

*Comparative analyses of the chigger microbiome*

The OTU table was transformed to a bacterial taxonomy table with raw read counts of each OTU presented in columns and samples in rows. The read count data were normalized to relative abundance, using the total sum scaling method or proportion. Bacterial communities were summarized with regard to sample groups (individuals and pools), selected chigger species and studied sites (mixed species), as well as soil samples from Thailand and Lao PDR. The dominant OTUs (the OTUs that presented proportionally ≥10% within a sample) were plotted as stack-bar charts in Microsoft Excel; whereas the OTUs that showed <10% in a sample were combined together in "others".

To calculate α-diversity, the read count data in the OTU table was normalized by different rarefaction depths at 100, 1,000, 10,000 and 100,000 reads per sample in order to optimize the best value for data normalization prior to further comparative analyses. Rarefaction subsampling at 10,000 sequences depth was then applied, as it showed the highest recovery of samples (95.41%) and OTUs (82.11%) compared to other depths. The diversity of bacterial OTUs among different sample groups (individual chiggers, pooled chiggers and soil) and categories (chigger species, habitats and study sites) was determined through richness and diversity estimators available as the default in QIIME; *i.e.,* observed richness, chao1 non-parametric richness estimator, and whole-tree phylogenetic diversity (PD_whole_tree). Non-parametric Kruskal-Wallis tests with Bonferroni post-hoc comparisons were performed to compare the alpha-diversity of bacterial OTUs among the sample groups.

Similar to the α-diversity analysis, a rarefaction depth at 10,000 reads per sample was first applied in the data normalization step prior to β-diversity analysis of bacterial composition among the sample groups. At the level of individual and pooled chiggers, β-diversity of bacterial composition among different categories (chigger species, habitat and study site) was computed by QIIME defaulted unweighted and weighted UniFrac (Unique Fraction) phylogenetic-based measurement methods. These methods take phylogenetic distance (that is, the fraction of tree length between sets of bacterial taxa) into account in calculating β-diversity metrics between pairs of samples [48]. Principal coordinate analysis (PCoA) was used to transform complex multidimensional data in the metrics into simplified orthogonal axes, and plotted them to visualize the clustering patterns of bacterial composition in the samples. The nonparametric ANOSIM method (Analysis of Similarity) with 1,000 permutations was used to test whether the clustering pattern of bacterial composition in the samples was statistically significant.

Geobacillus *qPCR and Sanger sequencing*

A pair of PCR primers was designed aiming to amplify a 16S rRNA gene portion for the genus *Geobacillus* and related Firmicutes. Ten representative sequences of Bacillales and *Geobacillus* OTUs derived from MiSeq sequencing runs and additional sequences of *Bacillus* spp. (KC443093.1, AB501343.1, DQ207730.2, AF058766.1, HM470251.1, DQ906100.1 and AF233579.1) from the NCBI nucleotide database were aligned using ClustalW multiple alignments. The following primers, 16SGbF (GTCCGGAATTATTGGGCGTA) and 16SGbR (TACGCATTTCACCGCTACAC) were used in qPCR, amplifying a 150-bp DNA fragment. The full-length *Geobacillus* amplicon product was synthesized by Eurogentec Ltd. (Southampton, UK) as a single-stranded oligonucleotide and used as a standard control in the qPCR assay.

Individual, 25-pooled and 50-pooled chiggers, as well as water samples from the laboratory water bath (Grant Sub; Grant Instruments, Cambridge, UK) and Qiagen microbial DNA-free water (negative control), were used in the qPCR assay. DNA from chiggers and 10 µl of water bath samples were extracted using the DNeasy Blood & Tissue Kit (Qiagen). Serial dilutions of oligonucleotide standard control from $5 \times 10^6$ to $5 \times 10^{-1}$ copies were used on each plate. The qPCR was carried out in 20-µl reactions containing 1 µl of DNA template, 1× SensiMix SYBR (Bioline, London, UK), 0.2 µM each primer and microbial DNA-free water. The qPCR was run with 35 cycles as follows: initial denaturation at 95°C for 10 min; 35 cycles of 95°C for 15 sec, 55°C for 30 sec, 72°C for 15 sec; and finally melting curve analysis, from 50°C to 95°C, in 0.5°C increments. The qPCR assays were run in a MiniOpticon Real-Time PCR System (Bio-Rad, Hercules, CA, USA), and quantitative data analysis was performed by CFX Manager Software v.3.1 (Bio-Rad). An analysis of variance (ANOVA) with Tukey's HSD post-hoc correction was performed in order to test differences in 16S rRNA gene sequence copies among the sample types.

Amplicons from the qPCR assays were visualized by 1.2% agarose gel electrophoresis, incorporating SYBR Safe (Invitrogen). The PCR products were then purified using the QIAquick PCR Purification Kit (Qiagen) and subjected to cloning using the pGEM-T Easy Vector System (Promega, Fitchburg, WI, USA). Ten recombinant *E. coli* JM109 colonies of each sample were inoculated in LB ampicillin broth and grown overnight at 37°C in a shaking incubator at 200 rpm. Plasmid DNA was extracted from the transformed-cell pellets using the Wizard Plus SV Minipreps DNA Purification Kit (Promega) following the manufacturer's protocol. Finally, the plasmid DNA samples were sent for Sanger sequencing with pUC/M13 forward and reverse primers to Source Bioscience (Nottingham, UK).

Bacterial taxonomy was assigned to the DNA sequences using RDP Naive Bayesian rRNA Classifier Version 2.10 [49] on the website https://rdp.cme.msu.edu. The taxonomical hierarchy was accepted at a >80% confidence threshold [50]. The DNA sequences were then aligned using ClustalW and phylogenetic tree construction was performed with the Maximum likelihood method using Mega software version 6.06 [51].

*Determination of GC content in 16S rRNA sequences*

We evaluated whether the influence of GC content differentially affected data obtained from individual and pooled chiggers (low and high DNA concentration templates, respectively). Representative sequences of the dominant bacterial OTUs from both individual and pooled chiggers were assessed for GC content using "Oligo Calc", an oligonucleotide properties calculator available at http://biotools.nubic.northwestern.edu/OligoCalc.html [52]. The mean GC content of the dominant OTUs were compared between individual and pooled chiggers by a parametric two-sample *t*-test.
